# Unified Transcriptome and Mechanics Map of the Intact Mammalian Preimplantation Embryo In Situ

**DOI:** 10.64898/2026.02.23.706394

**Authors:** Ehsan Habibi, Anubhav Sinha, Haiqian Yang, Payman Yadollahpour, Raquel Rodriguez-Aller, Yiwei Li, Lani Lee, David A. Wollensak, Zachary D. Chiang, Denny Sakkas, Edward S. Boyden, Armen Saghatelyan, Ming Guo, Aviv Regev, Fei Chen

## Abstract

The development and maintenance of multicellular tissues require that cell states be closely coupled to their local environment, including geometric and mechanical cues. However, studying this coupling in intact tissues has been challenging, because existing measurement technologies cannot simultaneously assess mechanical properties and molecularly defined cell states. To address this gap, we introduce the Unified Transcriptome and Mechanics Map (UTMM), a method for concurrent measurement of the transcriptome and cytoplasmic stiffness (high-frequency intracellular mechanical compliance) within intact 3D multicellular structures. UTMM relies on two innovations: (1) a targeted in situ RNA sequencing approach for intact 3D embryos (3DISS), and (2) a strategy that leverages high-frequency intracellular organelle fluctuations to provide a relative, non-invasive readout of cell-level viscoelastic compliance in situ. We applied UTMM to mouse embryos from the zygote through the morula stage to characterize how RNA expression and intracellular mechanical state becomes coupled during early lineage specification. Our data reveal that, in the early morula, transcriptional and morphological distinctions emerge between trophectoderm and inner cell mass (ICM) lineages, coinciding with a gradual increase in intracellular compliance (softening) across all cells from the 2-cell through morula stages. Furthermore, we observe that early lineage biases associated with distinct mechanical states, reflecting emerging developmental programs. When we osmotically delayed this softening, embryonic progression was impeded, supporting a functional relationship between intracellular material state and developmental timing. Extending this approach to neural stem cells in the adult brain (NSCs), we further show that proliferated NSCs exhibit greater intracellular compliance than quiescent NSCs, NSC state-dependent differences in the mechanical properties and that the analytical framework can resolve biologically associated mechanical differences in a distinct tissue context. Together, these results demonstrate UTMM’s ability to bridge molecular and mechanical dynamics in multicellular systems, providing a powerful framework for investigating how molecular state, geometry, and intracellular mechanics become coordinated during multicellular development.

**One Sentence Summary:** Joint 3D in situ quantification of spatial transcriptome and cytoplasm mechanics in preimplantation mammalian embryos

## Introduction

The development and maintenance of multi-cellular tissues is an intricate process that integrates genetic and molecular pathways with additional factors, including cellular geometry and mechanical cues ^1–5^. In particular, mechanical forces are critical in cell differentiation, tissue morphogenesis and organogenesis ^1,2^. Unraveling the interplay between mechanical properties and molecular pathways is a critical challenge in contemporary developmental and tissue biology. However, performing multi-modal measurements of both molecular profiles and mechanical properties of cells remains challenging, especially within intact 3D multi-cellular structures, such as embryos.

Recent technical advances are addressing some limitations, but substantial challenges remain. On the one hand, single-cell and spatial transcriptomics now allow us to measure gene expression profiles at both large scale and high cellular and spatial resolution. However, spatial transcriptomic approaches have been mostly restricted to thin (∼10 microns) 2D sections, and have not been applied to 3D intact embryos (∼100-150 microns) ^6^. On the other hand, despite important advances in probing the mechanical cues that dictate cell behavior in multicellular structures, existing techniques are often invasive and can only examine cells on the periphery of a 3D structure ^7–16^. Finally, no studies to date have combined both approaches at the needed scale to allow simultaneous measurements.

Mammalian preimplantation development is a prime example of self-organized morphogenesis. The transition from the single-cell zygote to the blastocyst is characterized by dynamic cell positioning and differentiation, with an intricate interplay between genetic programming, mechanics, and geometry, guiding cell behavior and physical arrangement. The development of the mammalian preimplantation embryo starts to progress through several steps from the single-cell zygote, with three rounds of cleavage giving rise to an 8-cell embryo, followed by compaction and polarization. During the morula stage, a tightly-packed assembly of 9 to 32 cells poised for cavitation, two pivotal cell fate decisions occur. The first lineage specification event segregates the trophectoderm (TE) from inner cells during the morula stage. Inner cells at this stage represent precursors to the Inner Cell Mass (ICM). The second lineage specification event, which segregates the primitive endoderm (PrE) from the epiblast (Epi) within the ICM, occurs later during blastocyst maturation ^17–19^. The TE is further subcategorized into the polar TE, in direct contact with the ICM, which eventually develops into the extraembryonic ectoderm (ExE) and ectoplacental cones, and the mural TE at the ab-embryonic portion of the blastocysts, which subsequently evolves into primary trophoblast giant cells ^20^. Although longstanding models posit that these early cell fate decisions are guided by both intrinsic gene regulatory circuits and extrinsic mechanical cues ^1–5^, it has been challenging to assess the contribution and mutual influence of such mechanisms, because of the difficulty to measure both molecular profiles and mechanical forces in intact 3D preimplantation embryos at scale.

Here, we introduce a novel approach for simultaneous quantification of expression profiles and mechanical properties at the single cell level *in situ*, and apply it to intact 3D mammalian preimplantation embryos to provide novel insights into early lineage specification events. First, we developed a targeted *in situ* RNA sequencing protocol for intact 3D specimens (3DISS) and used it to measure subcellular RNA expression and localization from the zygote to morula stage. Second, we developed a new approach to measure the mechanical properties of individual cells in live embryos. We computationally integrated these measurements into a Unified Transcriptome and Mechanical Map (UTMM): a multi-modal atlas of mechanical and expression dynamics in preimplantation embryos. Studying the UTMM, we characterize how mechanical properties, expression programs and physical architecture dynamically change during early lineage specification and subsequent differentiation and show how perturbing cellular mechanics patterns delay embryo development. Furthermore, we have also revealed intricate differences in the mechanical properties of NSCs at their activated and quiescent states. Our approach paves the way for a more comprehensive understanding of the complex interplay between mechanical and molecular signatures that underlie the first fate decisions of mammalian development.

## Results

### A framework for constructing a unified transcriptome and mechanical map (UTMM)

We developed a data collection and integration framework for constructing unified transcriptome and mechanical measurements in 3D tissue, such as embryos. To do so, we developed technical innovations along three data collection modules: (**1**) a novel approach for capturing cytoplasmic mechanics through passive microrheology, by measuring high-frequency fluctuations of organelles at single-cell resolution through live 3D imaging of whole embryos; (**2**) morphology measurements and automated segmentation algorithms to retrieve 3D geometrical maps of each cell within a preimplantation embryo; and (**3**) highly multiplexed spatial transcriptomics by *in situ* sequencing of intact 3D preimplantation embryos to capture molecular cell types and states. We collected each of these modules via microscopy in the same 3D embryo, followed by registration of the microscopy coordinate systems to align cytoplasmic mechanics with morphological and gene expression profile into a unified, multidimensional map for each embryo (**Fig. 1**, **Extended Data Fig. 1**). We describe each module and their integration in detail next.

**Figure 1.**
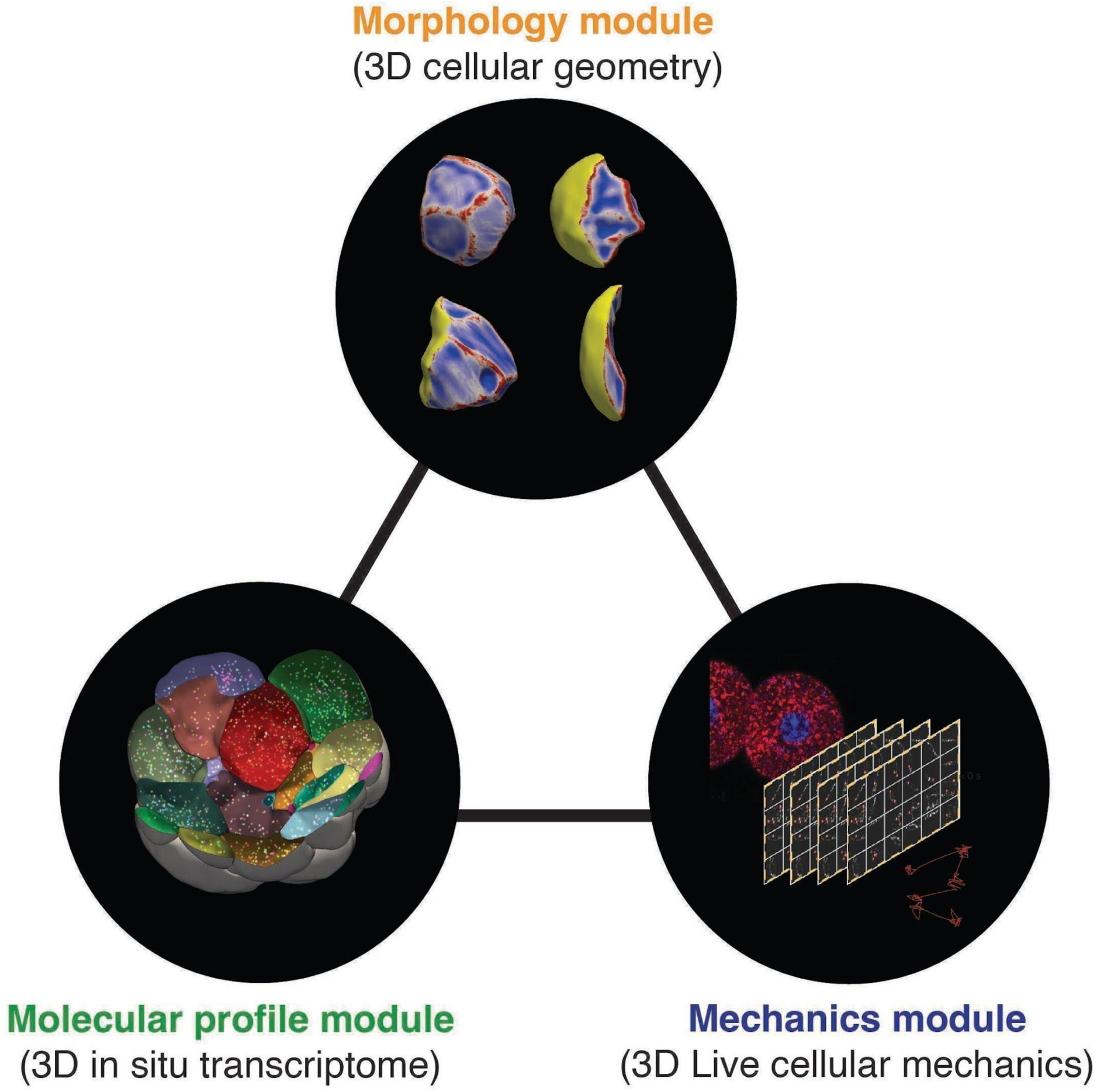
Unified Transcriptome and Mechanics Map (UTMM) modules. Overview of approach and data collection.

### Whole mount *in situ* RNA sequencing in preimplantation embryos

To measure gene expression dynamics across preimplantation development within intact embryos (**Fig. 2a**), we developed an approach for gel-embedding and clearing of fixed preimplantation embryos to allow multiple rounds of 3D confocal imaging (**Extended Data Fig. 1, Methods**), as well as for optimized multiplex antibody staining in the gel environment to enable robust 3D segmentation of individual cells (using E-cadherin, β-catenin and p-Ezrin) (**Fig. 2b**, i and iii and **Extended Data Fig. 2, Methods**). We modified an existing protocol for high sensitivity targeted *in situ* sequencing ^21,22^ for imaging and decoding the expression of 150 genes through *in situ* sequencing of barcoded probes in a 3D tissue (3DISS, **Methods**, **Supplementary Tables 1-3**). Specifically, we performed seven rounds of decoding across four color channels (**Fig. 2b**, ii). We optimized the *in situ* sequencing protocol for the thick 3D embryo, and validated that sequencing results are robust to Z depth (**Extended Data Fig. 3a-b).** We leveraged published scRNA-seq data of preimplantation embryogenesis to select a gene signature of expression changes from late 2-cell to late blastocyst stages ^23–25^. Using URD, we reconstructed a trajectory tree to model cell differentiation along pseudotime ^23–25^ (**Extended Data Fig. 4**), mapped expression changes of gene modules along each branch, identified top lineage markers, and selected 150 genes spanning different cell identities and cellular processes, including ligand-receptor pairs, morphogen signaling pathway genes, and transcription factors (**Supplementary Tables 1-3**, Supplementary Data 1, **Methods**).

**Figure 2.**
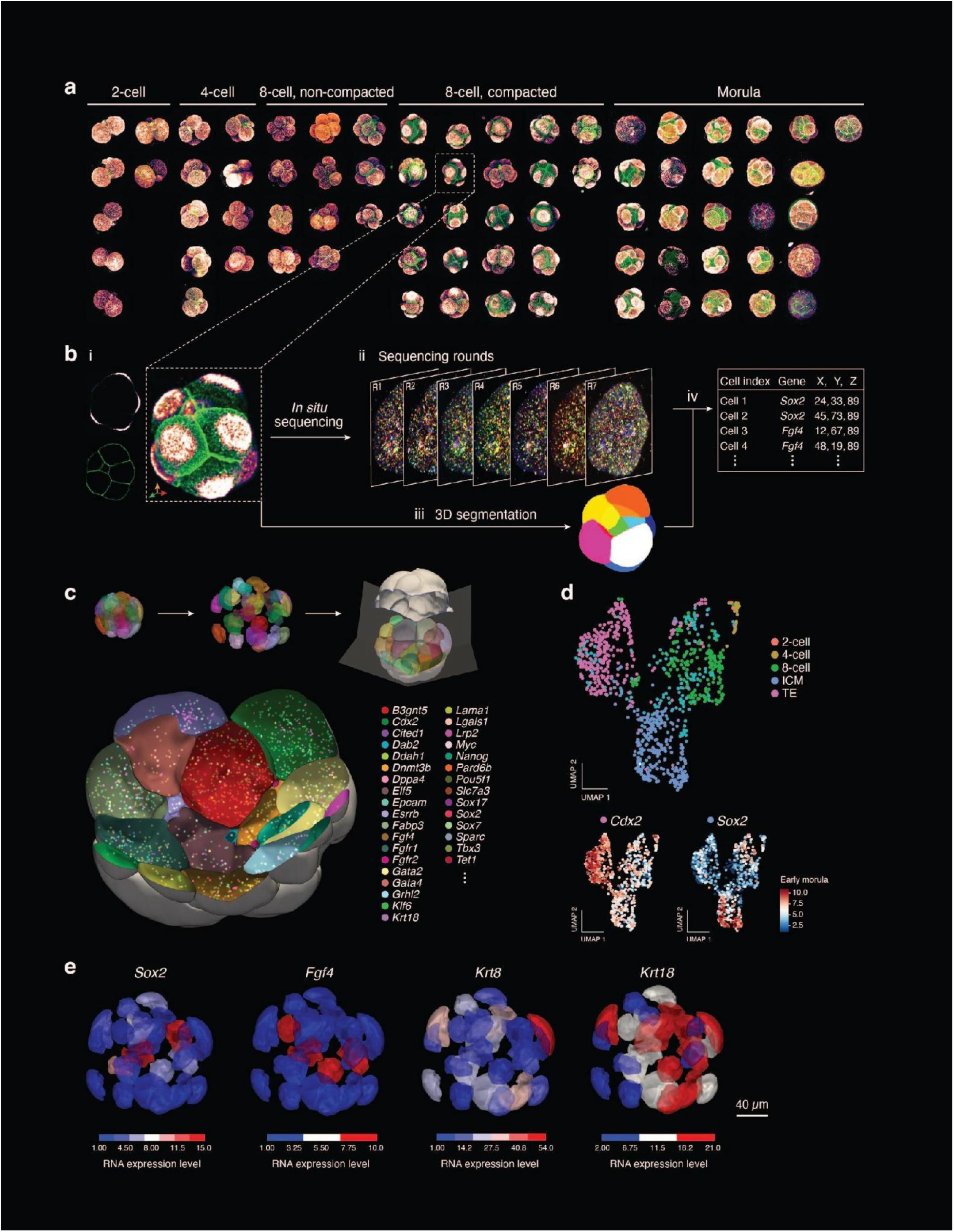
Spatiotemporal transcriptomics mapping in mouse preimplantation via whole mount targeted *in situ* sequencing. (**a**) 3D fluorescence microscopy images of 69 embryos captured across mouse preimplantation developmental stages. Colors depict antibody staining for E-cadherin and β-catenin (acquired together in the same color channel, green) and p-Ezrin (gem colormap). (**b**) Whole mount in situ workflow. (i) Embryos are fixed, embedded in gel, and subsequently immunostained (inset of A). (ii) Targeted in situ sequencing using RNA splinted padlock probes with gene specific barcodes (**Methods**). Shown are seven *in situ* sequencing rounds from an 8-cell stage embryo’s z-section. (iii) 3D segmentation of embryos from antibody staining. (iv) generation of spatial gene expression matrix for each cell within an embryo. (**c**) Transcripts sequenced within a 3D 16-cell stage embryo. Cells are enlarged to showcase individual RNA molecules (colored dots) from 33 distinct genes (out of 150 genes). (**d**) Distinct *in situ* cell profiles in different embryonic stages. Uniform manifold approximation and projection (UMAP) embedding of *in situ* RNA cell profiles (dots) colored by embryo stage (top) or expression levels of marker genes (bottom). (**e**) Distinct 3D expression patterns of canonical markers in a 16-cell stage embryo. 16 cell embryos images with cells colored by transcript count per cell of key marker genes. Scale bar: (E) 40 μm.

Next, we performed 3D targeted *in situ* sequencing of the 150 genes in 69 embryos spanning late 2-cell (*n*=6 embryos), 4-cell (*n=*7), 8-cell (*n=*32), and morula (*n=*24) stages, corresponding to E1.5 – E3.25 (**Fig. 2a**, **Methods**). We segmented the images, decoded the barcodes to identify individual mRNA molecules (**Fig 2b**; **Methods**), and assigned each decoded molecule to a cell (**Fig. 2b**, iv and **2c**, **Extended Data Fig. 5a**, ∼1,400 molecules/cell). In total, we profiled 742 cells across 69 embryos detecting 1,466,054 individual mRNA molecules (**Supplementary Table 4-5**). Our 3D ISS method effectively captured transcripts throughout the entire embryo, irrespective of cellular location, as demonstrated by comparable transcript detection in both internal and external cells (**Extended Data Fig. 3c**). Additionally, gene expression counts obtained through 3D ISS showed strong correlation with publicly available scRNA-seq^23–25^ datasets for corresponding embryo stages (**Pearson’s r = 0.55–0.80, Extended Data Fig. 3d**). Lastly, to assess the sensitivity of detection, we performed multiplex whole-mount *in situ* hybridization chain reaction (HCR) for 12 genes simultaneously on each individual embryo and compared the results with our 3D ISS. The comparison revealed that 3D ISS achieves an mRNA detection sensitivity approximately ∼64% of single-molecule FISH by HCR, with a strong correlation to HCR smFISH (Pearson’s r = 0.96, **Supplementary Table 6**).

### *In situ* cell states across key developmental transitions

The cell intrinsic profiles of all 742 cells measured *in situ* followed the main differentiation continuum from 2-cell to morula stage (**Fig. 2d**, **Methods**), such that the assigned pseudotime value of each cell (**Extended Data Fig. 5b**) aligned well with the true time point of sampling (**Fig. 2d**). Clusters were identified using unsupervised clustering and annotated into five main subsets based on stage and known markers (**Methods**): 2-cell, 4-cell, and 8-cell stages, and two morula subsets, one of TE-like cells and one of ICM-like cells (**Fig. 2d** and **Extended Data Fig. 6a**).

Key events in early development were associated with dynamic changes in cell intrinsic expression *in situ*. For example, genes upregulated in non-compacted *vs*. compacted 8-cell stage embryos (as identified via staging in **Fig. 2a**) included those pivotal for stem cell maintenance and *in utero* development (*Pou5f1*, *Tbx3*, *Essrb*, and *Rxra*) and others integral to cell junction assembly (*Cldn4*, *Cldn7*, and *Gja1*) (**Extended Data Fig. 6c**), consistent with the known morphological compaction and increased cell-to-cell contact in the compacted 8-cell stage ^26,27^. In the morula stage, the outer cell subset (TE-like) is characterized by *in situ* expression of *Cdx2*, *Krt8*, *Krt18*, *Fabp3*, and *Gata3*, and the inner subset (ICM-like) by genes such as *Sox2* and *Nanog* (**Fig. 2d** and **Extended Data Fig. 6a**, **Methods**), consistent with published scRNA-seq profiles (**Extended Data Fig. 6b**) ^23,28,29^.

Similarly, *in situ* expression patterns in a single morula stage embryo, when cells begin to diversify, accurately captured the expected spatial distribution of known markers. For example, pluripotency markers (e.g., *Sox2*, *Nanog*, and *Fgf4*) were predominantly expressed in cells near the center of the embryo, marking prospective ICM (**Fig. 2e** and **Extended Data Fig. 7** top); whereas markers of the prospective TE (*e.g.*, *Krt18*, *Krt8, Lrp2*) were more highly expressed in cells on the outer layer of the embryo (**Fig. 2e** and **Extended Data Fig. 7** bottom).

### Transcriptional divergence and spatial segregation co-emerge during the morula

Despite extensive research, the precise timing of cell fate decisions in the morula for epiblast (Epi), primitive endoderm (PrE), and polar and mural TE, remains incompletely elucidated ^28,30–36^. To identify the spatiotemporal expression patterns associated with the establishment of the ICM and TE, we stratified 24 morula-stage embryos into early morulae (9-16 cells; 11 embryos, 164 cells profiled *in situ*) and mid-late morulae (17-32 cells; 13 embryos, 278 cells profiled), learned gene programs from the profiles of the cells of each embryo set independently using non-negative matrix factorization (NMF) ^37,38^, annotated them by their associated genes, and related them to their spatial positions (**Fig. 3a**, **Extended Data Fig. 8a and b**, **Methods**).

**Figure 3.**
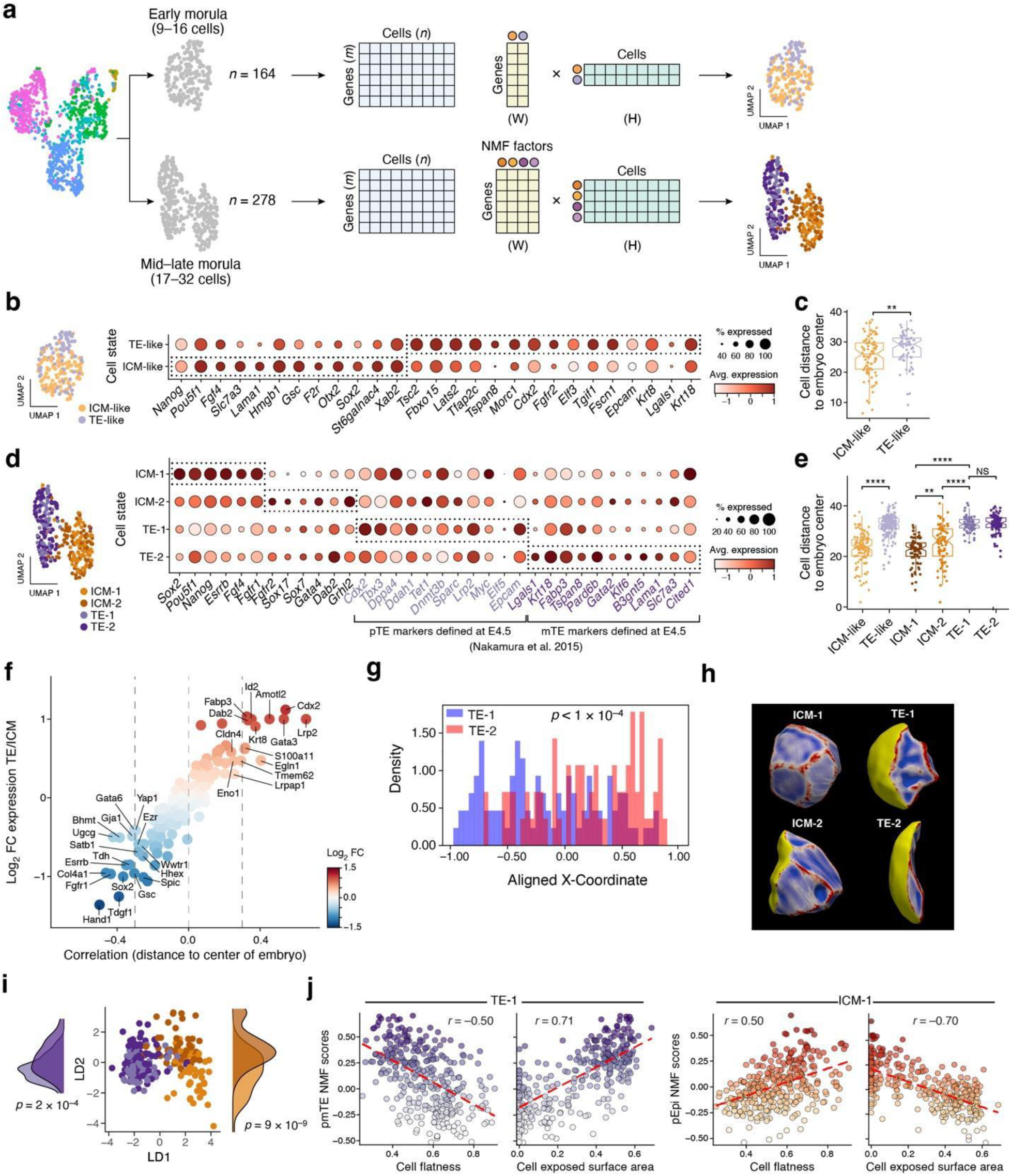
3D *in situ* sequencing of morula stage embryos reveals early ICM and TE lineage divergence. (**a**) Overview of non-negative matrix factorization (NMF) analysis. NMF programs are learned from cell profiles from early morula (9-16 cells, *k*=164) and mid-late morula (17-32 cells, *k*=278), and the top 15 genes for each program used to score cells. (**b-f**) *In situ* gene programs distinguish ICM and TE subsets with distinct spatial distributions. (**b,d**) Left: UMAP embedding of *in situ* cell profiles from early (b) and late (d) morula, colored by NMF-based annotation. Right: Mean expression (dot color; normalized transcripts count, colorbar) and fraction of expressing cells (dot size) of genes (columns) that are differentially expressed between different cell states (rows) in early (b) or late (d) morula. Purple: E4.5 mouse blastocyst genes from Nakamura et. al. 2015. (**c,e**) Distribution of Euclidean distances from cell center to embryo center (y axis) for cells from each state (x axis) in early (c) or late (e) morula. Boxplots denote the medians and the interquartile ranges (IQRs). Whiskers: lowest datum within 1.5 IQR of the lower quartile and the highest datum within 1.5 IQR of the upper quartile. Wilcoxon ranked sum test:, ns, p > 0.05 ; *, p ≤ 0.05 ; **, p ≤ 0.01 ; ***, p ≤ 0.001, ****, p ≤ 0.0001. (**f**) Correlation between expression level and mean radial position to the embryo’s center (x-axis; Pearson’s |r| > 0.3) for each gene (dot) differentially expressed between ICM and TE cells (y axis). Blue/red: higher expression in TE/ICM (p<10^-7^, Wilcox test). (**g-j**) Association between distinct expression and morphological features. (**g**) Spatial separation between TE-2 and TE-1 populations across embryos. Distribution of aggregated TE-2 (blue) and TE-1 (red) cells along the cross-embryo aligned X-coordinate axis (**Methods;** Kolmogorov-Smirnov test, p < 1 ×10^-4^). (**h**) Four example cells from a single mid-late stage morula embryo with cells colored by scores for cell surface area (blue), exposed surface area to the outside of embryo (yellow), and curvature (red). (**i**) Mid-late morula cell morphological profiles (dots) embedded in the space of the first two components of Linear Discriminant Analysis (LDA) (x and y axis) of 64 features colored by the cells’ expression based annotation.Marginal distributions: the density plot on the left shows the separation of TE subsets (TE-2 and TE-1) with a statistically significant difference (Wilcoxon test, p<2.0×10⁻⁴) and the density plot on the right depicts two populations within the ICM at mid-late morula (ICM-1 and ICM-2) also showing a significant separation (Wilcoxon test, p<9.0×10^-9^). (**j**) TE-2 (left panels) and ICM-1 (right panels) NMF scores (y axis) and proportion of exposed cell surface area or cell flatness (x axes) for cells (dot) from mid-morula stage. Dashed black line: best-fit line of linear regression between the variables. Top left: Pearson’s r.

Early morula cells had two main programs, annotated as TE-like and ICM-like based on the top-ranked 15 genes (**Extended Data Fig. 8a**), which we used to assign each cell as either ICM-like or TE-like (by spectral clustering using these factors on matrix **H**, **Methods**). The ICM-like and TE-like populations arrange along a continuum (**Fig. 3b**, left), with distinct expression patterns, including higher expression of *Sox2* and *Fgf4* in ICM cells, and of *Cdx2*, *Krt8*, and *Tfap2c* in TE cells (**Fig. 3b**) ^28,29^.

Correspondingly, the cells had different spatial distributions, as defined by the Euclidean distance from each cell to the center of the embryo: on average, ICM-like cells were closer to the embryonic center, and TE-like cells were more peripheral (Wilcoxon test, p < 0.01), yet considerable positional overlap existed at this early stage (**Fig. 3c**). Moreover, both the strength of the ICM-like signature and the difference between the ICM-like and TE-like signature of each cell were inversely correlated to the embryonic center (Pearson’s r = -0.33, -0.4, p <10^-4^), such that cells closer to the center have stronger ICM-like signatures and weaker TE-like signatures (**Extended Data Fig. 8c**, left). Thus, lineage biases emerge early at the RNA expression level and their magnitude is associated with cell positioning, but as considerable positional overlap persists, there is also inherent plasticity. Together with prior studies demonstrating that mechanical and morphological factors, such as polarity and cortical tension, influence fate decisions, these results might suggest that expression programs and spatial or mechanical cues co-emerge and interact to reinforce and stabilize lineage identity over time ^7,39,40^.

In the mid-late morula stage, the ICM-like (*Sox2*+, *Fgf4*+) and TE-like (*Cdx2*+, *Krt8*+) subsets have further separated in expression (compare **Fig. 3b** and **3d**, left) as well as physical space (compare **Fig. 3c** and **3e**). By mid-late morula there is a clearer spatial segregation of inner and outer embryo for molecularly annotated ICM and TE cells (Wilcoxon test, p < 10^-4^; **Fig. 3e,f**), and a strong negative correlation between a cell’s ICM program score or the difference between its ICM and TE scores and its distance from the embryo center (Pearson’s r = -0.60, -0.88, p < 2.2×10^-16^, **Extended Data Fig. 8c**, bottom). This is consistent with a model of ongoing concomitant divergence in both physical space and transcriptional identity: ICM cells intensify their ICM lineage-specific gene expression and reduce their TE related expression, while becoming increasingly centralized within the embryo, whereas TE cells demonstrate more peripheral positioning and expression identity (and reduce ICM related expression). Consistently, at the individual gene level, higher expression in TE cells (*e.g.*, *Cdx2*, *Krt8*, *Lrp2*, *Amotl2*, *Id2*, *Cldn4*, *Egln1*, *Fabp3*, *S100a11, Gata3*; **Extended Data Fig. 8e**) was positively correlated with distance from the embryo center (Pearson’s r > 0.3, p < 10^-7^, **Fig. 3f**), and higher expression in ICM cells (*Sox2*, *Esrrb*, *Gsc*, *Tdgf1*, *Spic*, *Tdh;* **Extended Data Fig. 8f**) was negatively correlated (Pearson’s r < -0.3, p<10^-7^, **Fig. 3f**).

### Early ICM and TE sub-subsets show spatial biases

Within the ICM and TE subsets in the mid-late morula stage, spectral clustering over the NMF programs resolved two early transcriptional sub-states in each group. In the ICM population, one sub-state showed higher relative expression of epiblast-associated genes (e.g. *Sox2*, *Fgf4*, *Pou5f1*, *Nanog*), while the while the other exhibited modest upregulation of primitive-endoderm-associated transcripts (*Fgfr2*, *Gata4*, *Sox7*, *Dab2*, *Sox17*)^28^(**Fig. 3d, Extended Data Fig. 8b**). We therefore refer to these as ICM-1 (“Epiblast-biased”) and ICM-2 (“PrE-biased”) sub-states, labels that reflect early transcriptional tendencies rather than fixed fates. To assess the robustness of these inferred programs, we repeated NMF decomposition on the inner-cell population across 1000 random initializations and quantified agreement using the adjusted Rand index (ARI), which measures similarity of cell assignments while correcting for chance. The resulting program assignments were highly reproducible (median ARI = 0.94, median ARI = 0.95), indicating stable identification of early transcriptional sub-states. Additionally, both ICM-1 and ICM-2 cells were more centrally positioned than TE cells (Wilcoxon test, p < 0.0001), with ICM-2 cells more outwardly biased than ICM-1 cells (Wilcoxon test, p < 0.01) (**Fig. 3e**).

Within the TE population, two sub-states could be distinguished by differential expression of known TE markers: one with higher relative *Cdx2*, *Gata3*, *and Ddh1* consistent with a “polar-biased TE” transcriptional tendency, and another with elevated *Krt18*, *Fabp3*, *and Gata2* expression, reflecting a “mural-biased TE” tendency (**Fig. 3d**, **Extended Data Fig. 8b**) ^41,42^. Accordingly, we refer to these TE subpopulations as TE-1 (“polar-biased TE”) and TE-2 (“mural-biased TE”) to emphasize their transcriptional heterogeneity and plasticity rather than imply a fully committed identity. Spatially, TE-2 and TE-1 cells were similarly positioned relative to the embryo center (Wilcoxon test, p > 0.05. **Fig. 3e**), but more peripheral than ICM (ICM-1 and ICM-2) cells (Wilcoxon test, p < 0.0001, **Fig. 3e**). Thus, TE-2 and TE-1 expression differences are apparent as early as the morula stage, prior to cavitation.

While cells in the TE-1 and TE-2 states are at a comparable distance from the embryonic center, we investigated whether they are spatially polarized in terms of opposite sides of the embryo. To address this, we developed a custom alignment method (**Methods**) that, for each embryo, identified a unit vector maximizing the separation of these two populations along a projection axis. This alignment ensured consistent relative positioning of TE-1 and TE-2 cells across embryos. By aggregating data into a unified coordinate system, we uncovered statistically significant spatial separation between TE-1 and TE-2 distributions (Kolmogorov-Smirnov test: D = 0.341, p = 1.1 × 10⁻⁴). These results reveal subtle yet reproducible spatial biases among transcriptionally biased TE subpopulations at the mid-to-late morula stage, suggesting that molecular priming and spatial organization may begin to emerge before lumen formation (**Fig. 3g**)

### Expression states are associated with distinct cell morphological features

Leveraging our high-resolution multiplex spatial transcriptome, which enables precise identification of early lineage-biased cellular states, we next asked whether these expression-defined populations correspond to reproducible differences in 3D cell morphology. Previous quantitative analyses of early mouse embryos demonstrated that positional geometry, rather than overall cell shape, predicts Hippo/YAP signaling and TE versus ICM fate bias^43^. However, whether similar geometric signatures distinguish transcriptionally defined lineage states has not been examined. To show this, we extracted 64 cell morphological, textural, and intensity-based features from 3D segmented cell shapes^44^ (**Fig. 3h**, **Methods**), and performed Linear Discriminant Analysis (LDA)^45^ on these feature profiles with two components as predictors and four previously-defined cell states as response variables (ICM-1, ICM-2, TE-1 and TE-2), to recover a feature space that maximizes the separation between the different expression cell states (**Fig. 3i**). The first component (LD1) separated the cells into two main clusters, TE and ICM, with additional distinction of ICM-2 from ICM-1 (**Fig. 3i**, Wilcoxon test, p<9.0×10^-9^) and TE-1 and TE-2 (**Fig. 3h**, Wilcoxon test, p<2.0×10^-4^) on the second component (LD2), suggesting that the molecular profiles have corresponding morphological distinctions (**Fig. 3i**, **Extended Data Fig. 9**). For example, cell scores for the TE-2 programs were negatively correlated with cell flatness (Pearson’s r = -0.51, p<2.2×10^-16^, **Fig. 3j**) and positively correlated with the proportion of exposed cell surface area (r = 0.7, p<2.2×10^-16^, **Fig. 3j**), and conversely for the ICM-1 program (r = 0.5 for cell flatness; r= -0.7 for exposed cell surface both p<2.2×10^-16^, **Fig. 3j**).

Early ICM cells can either form internally or migrate inward from the outer layer. To investigate the spatial behavior of transcriptionally defined ICM-derived cells and assess what proportion appears on the outer layer before potentially migrating inward, we analyzed the exposed cell surface area normalized to total cell surface area. Interestingly, we observed a continuum of exposed surface area values for ICM-1 and ICM-2 cells (**Extended Data Fig. 9**). This morphological continuum was further validated by visually inspecting 3D-segmented embryos (**Extended Data Fig. 10**), where cells with minimal exposure (<0.1) represented either truly internal cells or potential segmentation artifacts, while cells with higher exposure (>0.15) reflected the transitional nature of cells during early ICM development. In the ICM-1 group, 70% of cells exhibited an exposed surface area below 0.05, 85% below 0.15, and none exceeded 0.25, indicating that ICM-1 cells predominantly reside internally, with a small fraction possibly undergoing inward migration. In contrast, ICM-2 cells displayed a higher degree of surface exposure, with only 40% having an exposed surface area below 0.05, 60% less than 0.15, and 35% exceeding 0.25. This suggests that ICM-2 cells, while some are generated internally, others originate on the outer layer and are observed in transitional phases as they migrate inward or undergo internalization, retaining partial surface exposure. This is consistent with previous studies showing that PrE-biased cells can originate at the embryo surface and subsequently undergo inward repositioning/internalization during morula-to-blastocyst transitions^30,31,46^. These findings indicate that transcriptional priming of ICM-derived cells, particularly the **ICM-2** program, can be detected while cells remain externally positioned, suggesting that lineage-biased gene expression arises alongside cell repositioning rather than strictly after internalization. Importantly, these observations do not imply that lineage fate is fixed or deterministically predictable at the morula stage. Rather, our data reveal incipient transcriptional biases that coexist with the well-established plasticity of early blastomeres. Cells remain capable of lineage switching through the 16–32-cell stage and beyond, consistent with classical lineage-tracing studies. Our measurements therefore identify early molecular asymmetries, not irreversible fate commitment.

### 3D *in situ* measurement of cytoplasmic stiffness based on endogenous organelle motion

Given the relationship between a cell’s expression state, position, and morphology within the embryo, we next sought to assess the relationship of these features to the cell’s properties that determine their response to mechanical forces, in particular cytoplasmic stiffness and viscoelasticity. While several methods exist for assessing such mechanical properties, their application is often hindered by their invasive nature and inability to probe cells within a 3D tissue ^47^. One compelling approach for non-disruptive measurement of mechanical properties would be passive microrheology, a technique to measure the viscoelastic properties of soft materials ^48–51^. Although passive microrheology in living cells is often influenced over longer timescales by non-equilibrium forces from active processes like motor proteins, obscuring intrinsic mechanical properties, we have previously shown that at shorter timescales (high frequencies), thermal fluctuations dominate, approximating thermodynamic equilibrium ^50,51,50–52^ ^52^. Therefore, high-frequency passive microrheology provides reliable measurements of cytoplasmic stiffness.

Conventionally, passive microrheology studies track spontaneous movement inside the cytoplasm using endocytosed microbeads (0.5-1 μm in diameter), but such beads have restricted cellular uptake in 3D tissue and may be cytotoxic during prolonged culture^53^. To overcome these limitations and minimize interference with the embryos, we developed a refined, bead-free approach for mechanical measurements based on the endogenous spontaneous motion of organelles, such as mitochondria or lysosomes (**Fig. 4a,b**). We reasoned that, as these organelles are naturally present and uniformly distributed in the cytoplasm with an approximate diameter of 1 μm, they could serve as ideal, non-invasive alternatives to microbeads for characterizing the physical environment inside living cells ^51,54,55^.

**Figure 4.**
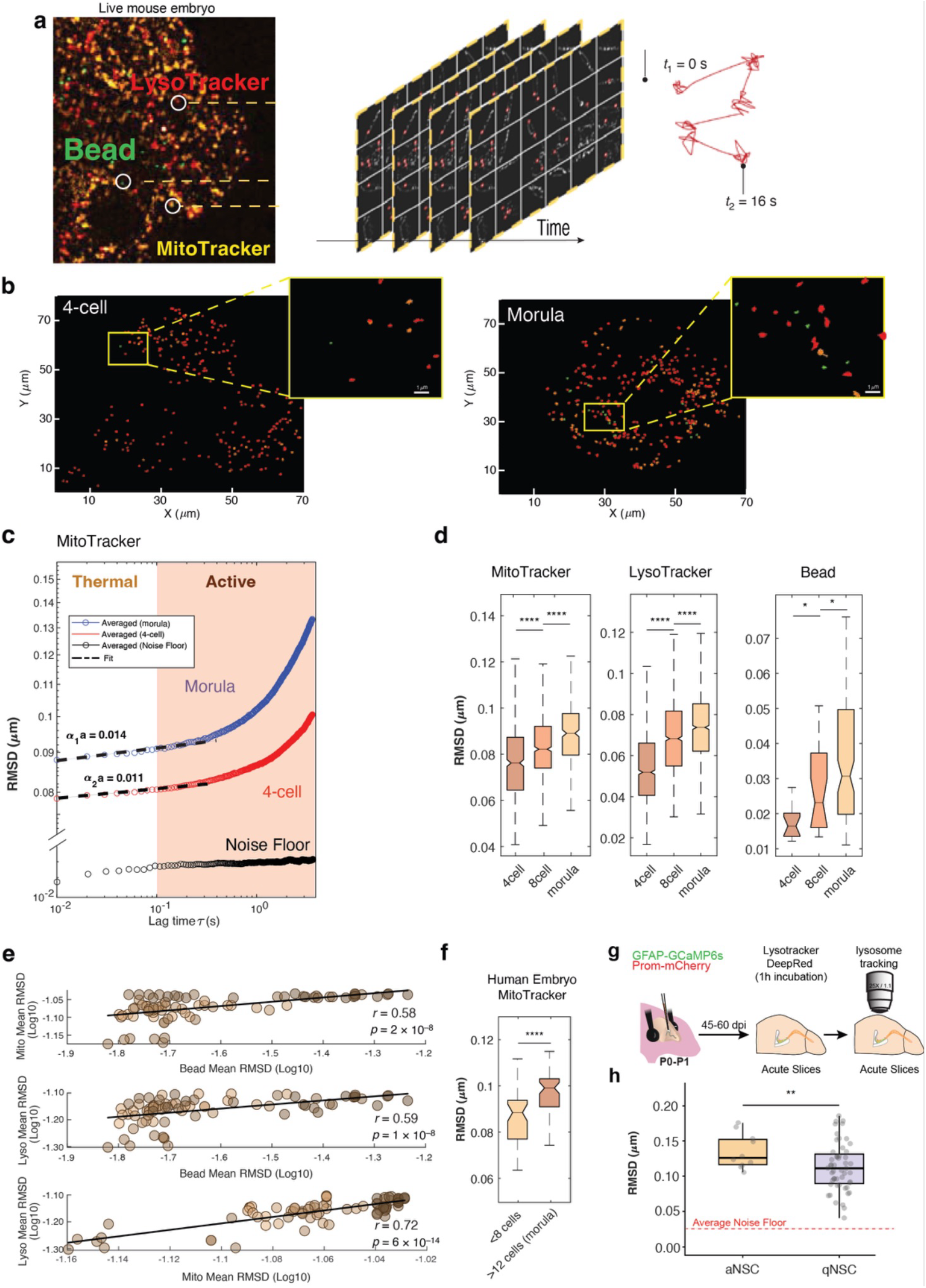
Spatiotemporal mapping of cytoplasmic mechanics in mammalian preimplantation embryos and mouse brain. (**a**) Embryo staining and imaging for particle tracking. Left: Single z-section of embryos co-stained with MitoTracker™ Red CMXRos (yellow, mitochondria), LysoTracker™ DeepRed (red, lysosomes) and endocytosed fluorescent microbeads (green) from confocal z-stack, acquired on a spinning-scanning confocal microscope. Middle: Selected fields of view from time-lapse videos (frame rate: 10 ms/frame) of each cell within an embryo. Red circles: mitochondria trajectories. Right: A schematic trajectory of an individual mitochondrion highlighting both short-term fluctuations in equilibrium and long-term active transport. Scale bar: 10 μm. **(b)** Example particle trajectories. Example trajectories for Mitochondria (yellow), lysosomes (red) and beads (green) at 4-cell stage (left) and morula stage (right). The trajectories were captured at a frame rate of 100 Hz for a total duration of 16.6 seconds (1667 frames). (**c**) Distinct mechanical regimes at different time scales. Root mean-squared displacement (RMSD, y axis, log scale) of fluorescently labeled mitochondria, measured with the particle tracking method, and lag time (x axis, log scale; the time interval over which the displacement of mitochondria is measured) for 4 cell (red) and morula (blue). Black line: noise level, measured by imaging and tracking the fluorescently labeled mitochondria in the same embryo after fixation. The logarithmic slope at high frequency is 0.011 for the 4-cell stage and 0.014 for the Morula stage. (**d**) High-frequency cytoplasmic motion increases during early embryonic progression. Distribution of RMSD values (y axis) recorded at each stage (x axis) using MitoTracker (left), beads (middle), or LysoTracker (right). Boxplots denote medians and IQRs. Whiskers: lowest/highest datum within 1.5 IQR of the lower/upper quartile. One-way ANOVA with Tukey’s multiple comparisons test, ns, p > 0.05 ; *, p ≤ 0.05 ; **, p ≤ 0.01 ; ***, p ≤ 0.001, ****, p ≤ 0.0001. (**e**) Correlation of mechanical properties between bead, mitochondrial, and lysosomal dynamics. Log-transformed Mean RMSD values for beads (x axis, top) and their 20 nearest mitochondria (y axis, top; Spearman’s ρ = 0.58, p = 2 ×10^-8^), beads (x axis, middle) and their 20 nearest lysosomes (y axis, middle; ρ = 0.59, p = 2 ×10^-8^), and mitochondria (x axis, bottom) and their 20 nearest lysosomes (y axis, bottom; ρ = 0.72, p = 6 ×10^-14^). Black lines: linear fits. (**f**) High-frequency cytoplasmic motion increases along human early embryonic development. Distribution of RMSD values (y axis) recorded before (N=2, 4-6 cell stage) and during (N=2, 14-16 cell) the morula stage (x axis) using MitoTracker. Boxplots denote medians and IQRs. Whiskers: lowest/highest datum within 1.5 IQR of the lower/upper quartile. One-way ANOVA with Tukey’s multiple comparisons test, ns, p > 0.05 ; *, p ≤ 0.05 ; **, p ≤ 0.01 ; ***, p ≤ 0.001, ****, p ≤ 0.0001. (**g**) Schematic representation of the experimental approach used to label adult neural stem cells (NSCs) by electroporation and assess intracellular stiffness from lysosome dynamics. (**h**) Distribution of RMSD values measured using LysoTracker in activated NSCs (aNSCs) and quiescent NSCs (qNSCs). Box plots show medians and interquartile ranges (IQRs); whiskers extend to the lowest and highest values within 1.5 × IQR of the lower and upper quartiles, respectively. One-way ANOVA with Tukey’s multiple-comparisons test; ns, P > 0.05; *, P ≤ 0.05; **, P ≤ 0.01; ***, P ≤ 0.001; ****, P ≤ 0.0001.

To measure high-frequency stiffness at single-cell resolution in live embryos, we stained each of 50 embryos across stages (4-cell, 8-cell and morula) with a live-cell compatible mitochondrial dye (**Fig. 4a-c**, MitoTracker™ Red CMXRos, **Methods**), and estimated spontaneous cytoplasmic motion by calculating the root mean squared displacement (RMSD) of spherical mitochondria. The RMSD_0_ curves reveal two distinct regimes (**Fig. 4c**). At shorter timescales (τ < 0.1 s), the RMSD_0_ reaches a high-frequency plateau, indicating thermodynamic equilibrium, where the cytoplasm behaves like a weak-elastic solid. This plateau value, RMSD0, serves as an indicator of mechanical compliance, with higher RMSD_0_ values reflecting a more compliant (less stiff) cytoplasm. The logarithmic slope (α) in this region is approximately ∼0.01, consistent with sub-diffusive behavior and a weak elastic response^51^. At longer timescales (τ > 0.1 s), the RMSD increases linearly, with slopes (α) reflecting motor-driven, diffusive behavior due to active transport mechanisms. This high-frequency analysis allows us to assess stiffness without interference from these active processes, providing a reliable measure of intrinsic cytoplasmic stiffness.

RMSD_0_ values are lower at the 4-cell stage compared to the morula, indicating that the cytoplasm is inherently stiffer at earlier stages (**Fig. 4c and d**). This suggests a more constrained cytoplasmic environment at the 4-cell stage, which may reflect developmental requirements for stability and structure in early embryo organization. Notably, even at these short timescales, the RMSD_0_ fluctuations in the stiffest stage (4-cell) were at least 7 times larger than the noise floor, ensuring reliable measurements above background noise (**Fig. 4c**). The RMSD_0_ values derived from mitochondria tracking gradually increase as the embryos develop, indicating a progressive decrease in cytoplasmic stiffness, or softening (**Fig. 4d**, one-way ANOVA, p < 2.2×10^-16^, with Tukey’s multiple comparisons test).

To validate mitochondria as a proxy for measuring cytoplasmic stiffness and determine whether other organelles could serve similarly, we conducted a parallel experiment to jointly quantify spontaneous movement using endocytosed microbeads (fluorescent carboxylate-modified polystyrene tracer particles, 0.5 μm in diameter), and another endogenous organelle (lysosome) (**Fig. 4a,b, Methods, Supplementary Video 1-5**). After dissolving the zona pellucida at the two-cell stage, we cultured the embryos with microbeads through the morula stage and co-stained with MitoTracker™ Red CMXRos and LysoTracker™ Red and imaged the same field of view across different stages to directly compare the motion of beads, mitochondria and lysosomes. Remarkably, RMSD_0_ values from beads, mitochondria, and lysosomes exhibited a highly consistent trend across developmental stages (**Fig 4d**). All were highly confined (stiffer) at the 4-cell stage but showed significantly greater dynamics at later stages (8-cell and morula), reflecting a consistent trend of cytoplasmic softening over time.

Next, to further assess local cytoplasmic mechanics and the relation between spontaneous motions of beads, mitochondria and lysosomes, we calculated mean RMSD_0_ values for mitochondria and lysosomes with respect to a 20-nearest-neighbor (20-NN) analysis centered around each bead (**Fig. 4e**). A moderately strong positive correlation was observed between bead RMSD_0_ and mitochondrial RMSD_0_ (r = 0.58, p < 0.0001) or lysosomal RMSD_0_ (r = 0.59, p < 0.0001) in its 20-NN proximity, indicating that mitochondrial or lysosomal movement closely reflects the local cytoplasmic dynamics near the bead. A stronger positive correlation between mitochondrial and lysosomal RMSD_0_ values (r = 0.72, p < 0.0001) further supports the close association between these organelles’ movements, reinforcing their use as non-invasive indicators of cytoplasmic stiffness.

To further confirm this, we also performed active microrheology using optical tweezers to sinusoidally oscillate these endocytosed particles at a frequency of 1 Hz and a small amplitude of 0.2 µm. The average magnitude of shear modulus of the cytoplasm of these embryos dropped from ∼35 Pa at the 2-cell and 4-cell stages to ∼10 Pa at the 8-cell stage, and even lower at the 16-cell stage (**Extended Data Fig. 11**). Overall, both optical tweezers active microrheology and high-frequency mitochondria movement consistently show gradual cell softening during preimplantation embryo development.

Additionally, we investigated whether the observed trend of cytoplasmic softening during cleavage stages is conserved in human preimplantation embryos. Using MitoTracker-stained human embryos, we measured mean RMSD_0_ values at different stages. As shown, RMSD_0_ values significantly increase from the <8-cell stage to the morula stage (>12 cells), indicating a consistent pattern of cytoplasmic softening similar to that observed in mouse embryos (**Fig 4f**). This suggests that the mechanical properties linked to developmental progression may be conserved across species.

Measuring intracellular mechanical properties within intact tissues, where cells reside in their true native microenvironment, remains a key unsolved challenge. To test whether organelle-based passive microrheology could meet this challenge in a fundamentally different biological context, we extended our approach to adult neural stem cells (NSCs) located in the sub-ventricular zone (SVZ) bordering the lateral ventricles of the adult forebrain. We asked whether intracellular stiffness could distinguish the two functional states of NSCs, activated (aNSCs) and quiescent (qNSCs), whose balance governs lifelong neurogenesis. Adult mice were co-electroporated at P0–P1 with GFAP-GCaMP6s and Prom-mCherry plasmids, exploiting the coincident activity of GFAP and Prominin promoters to specifically dual-label NSCs (**Fig. 4g** and **Extended Data Fig. 12**)^56–58^. Lysosome dynamics were tracked using LysoTracker DeepRed in 250-μm acute sagittal brain sections at 74.6 frames/s (**Extended Data Fig. 12a,b**). Post-hoc Ki67 immunolabeling was used to distinguish activated NSCs (aNSCs; Ki67+) from quiescent NSCs (qNSCs; Ki67−). We found that aNSCs exhibited significantly higher lysosome displacement compared to qNSCs (**Fig. 4h**), consistent with a softer cytoplasm in the activated state. These findings demonstrate that lysosome-based passive microrheology can resolve biologically relevant differences in intracellular mechanics across distinct cell states in the adult brain tissue, further demonstrating the generality of this approach.

Thus, high-frequency organelle movements such as mitochondria and lysosomes can be used as an indicator of cytoplasmic compliance for non-invasive monitoring of spatiotemporal dynamics of mechanical properties within intact 3D tissues, from early embryogenesis to cells located deep within adult brain tissue, establishing a broadly applicable tool for probing intracellular mechanics across tissue types and biological contexts.

### Unified transcriptome and mechanics show association between lineage and cell softness

Next, we examined the relationship between spatial position, expression profile and mechanical properties of each cell within individual embryos, by a dedicated experimental and computational workflow (**Fig. 5a**, **Methods**). Briefly, at each embryo stage, we first scanned through all the cells in the embryo with high-frequency videos (to perform passive microrheology), recorded the microscope stage positions for each video, scanned the entire embryo to produce a z-stack image for live-stained nuclei, and then fixed, immunostained and *in situ* sequenced the embryo (**Methods**). We computationally merged the mechanics and expression data for each cell and embryo, by registering the nucleus images collected during live-cell imaging for cytoplasmic mechanics with those from fixed imaging for RNA *in situ* sequencing, by manual 3D registration followed by affine registration (**Fig. 5a**, **Methods**). In total, 69 embryos were profiled by *in situ* RNA sequencing, 55 embryos were subjected to passive microrheology, and a subset of 29 embryos (438 cells) were jointly measured by both and registered with our pipeline.

**Figure 5.**
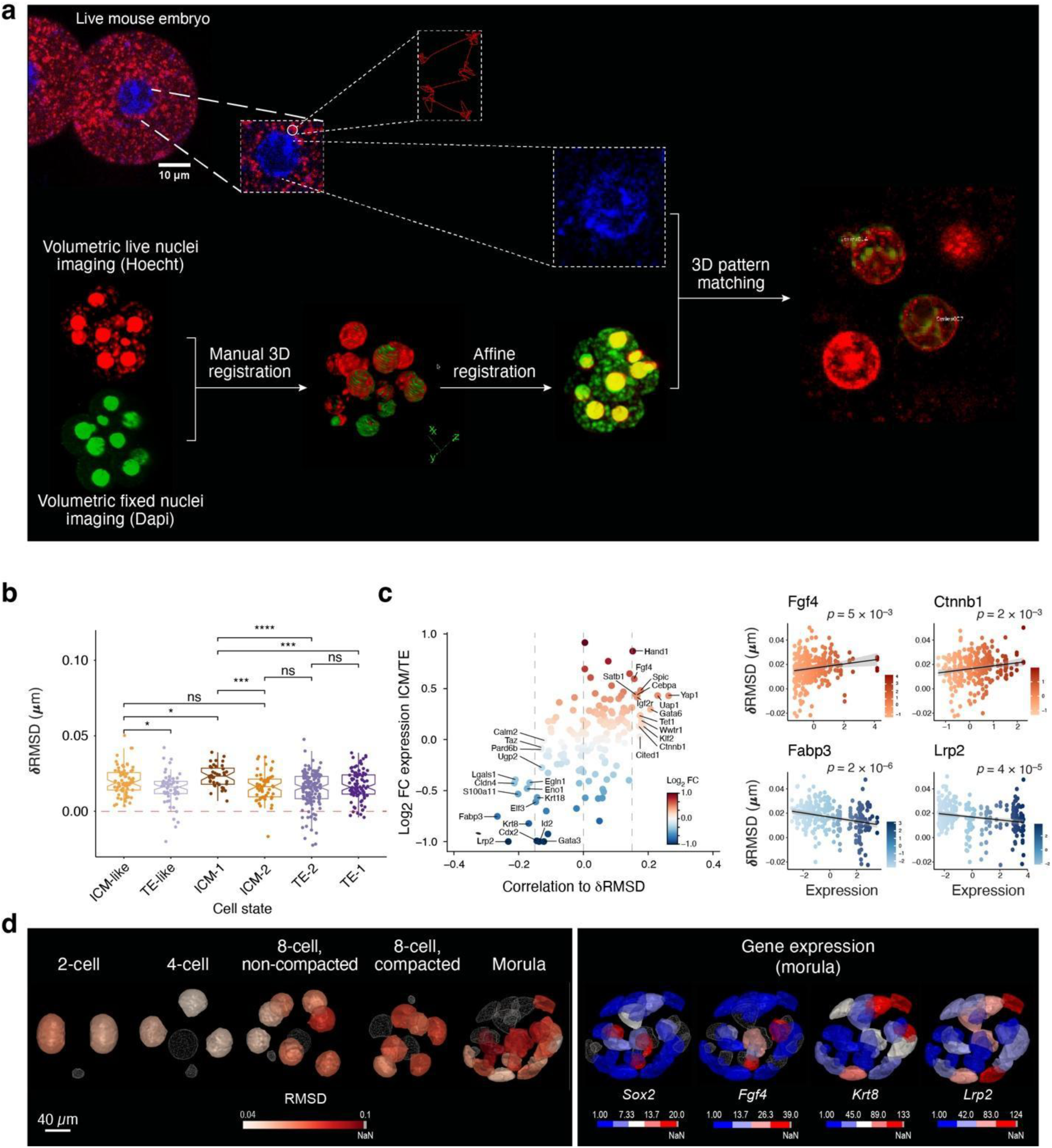
Workflow for computational mapping of mechanical and transcriptome measurements. (**a**) Left: Volumetric nucleus images from live (for cytoplasmic mechanics) and fixed (immunofluorescence and RNA *in situ* sequencing) embryos are manually registered in 3D bridges using volumetric nuclei images, followed by 3D affine registration. Top and right: All single frame nuclei (Hoechst) from particle tracking are pattern matched against volumetric DAPI (reference) to locate corresponding cells to pair each cell with both expression and RMSD data. (**b**) Distinct mechanical properties of ICM and TE biased cells in early morula. Distribution of δRMSD (y axis) in ICM and TE cell subsets (x axis). The red dashed line (at y = 0) indicates the 4-cell as a reference. Boxplots denote medians and IQRs. Whiskers: lowest/highest datum within 1.5 IQR of the lower/upper quartile. ns, p > 0.05 ; *, p ≤ 0.05 ; **, p ≤ 0.01 ; ***, p ≤ 0.001, ****, p ≤ 0.0001, Wilcoxon ranked sum test. (**c,d**) Gene expression levels correlated to mechanical properties. (**c**) Left: Correlation between expression level and RMSD (x-axis) for each gene (dot) across cells and its fold change between ICM and TE cells (y-axis). Blue/ red higher expression in TE/ICM. Only genes with Pearson’s |r| > 0.15 and p<0.01 [Wilcoxon Rank Sum test] are shown. Right: Expression (x axis) and RMSD (y axis) in each cell from morula stage embryos for different genes (label on top). (**d**) Left: RMSD values of cells from illustrative 2-cell to morula stage embryo. Right: Expression (color bar) in each cell of selected canonical markers the morula stage embryo also presented on left. Scale bar: 40 μm.

During the early morula window (9-16 cells) ICM-like cells were softer on average (higher δRMSD values) than TE-like cells (p < 0.05, Wilcoxon test, **Fig. 5b**). δRMSD, defined as the deviation of RMSD from the 4-cell as the earliest stage, captures relative differences in cytoplasmic softness across developmental stages. By the mid-late morula stage (16-32 cells), ICM-1 cells were on average softer than early ICM cells (p < 0.05, Wilcoxon test, **fig. 5b**), whereas ICM-2 cells remained stiffer than ICM-1 cells (p < 0.0001, Wilcoxon test, **Fig. 5b**), and comparable to early ICM, TE-2 and TE-1 cells (**Fig. 5b**). Consistently, genes whose expression was positively correlated with higher δRMSD were associated with pluripotency and ICM-specific markers (Pearson’s r > 0.15, *e.g.*, *Fgf4, p=*5×10^-3^ and *Ctnnb1, p=*2×10^-2^; **Fig. 5c and 5d**) and those negatively correlated associated with the TE lineage (Pearson’s r < -0.15, *e.g.*, *Fabp3, p=2*×10^-6^ and *Lrp2, p=2*×10^-5^; **Fig. 5c**). Thus, lineage segregation is associated with coordinated changes in cell morphology, expression profile, and softness.

### Volumetric compression delays embryo development

Recent studies suggest that molecular (de-)crowding might be a switch for certain signaling pathways ^59,60^ and that high-frequency stiffness may reflect cytoplasmic crowdedness ^61,62^. Thus, we examined whether developmental progression can be tuned through volumetric compression that delays a de-crowding (and hence softening) process. To investigate this, we cultured the embryos in a medium supplemented with 2% PEG 300 (400 mOsm), a polymer solution with a much higher osmotic pressure than the cytoplasm (200-250 mOsm), to osmotically force water efflux and increase the degree of molecular crowding in cells ^49,59^. We first grouped embryos that reached the 4-cell stage (**Fig. 6a**, **Methods**), cultivated them in either control (KSOM) media or supplemented with 2% PEG 300 (**Fig. 6a**, n = 200 embryos per group, 5 replicates, 40 embryos per replicate), and counted the number of embryos assigned to each stage at E2.5, E3.5, E4.5, and E5.5 days post-fertilization (**Fig. 6b**). Remarkably, volumetric compression delayed embryo development (**Fig. 6b**). For example, at E2.5, >80% of control embryos had advanced to the compacted 8-cell stage (early 8-cell = 2 embryos ± 0.63 (mean ± s.d.); late 8-cell = 32.80 ± 0.8 out of 40 in the replicate), but only 20% of the osmotically compressed group did so (early 8-cell = 28.6 ±3.12; late 8-cell = 7.20± 2.2). At E3.5, the developmental delay persisted in the osmotically compressed embryos (morula = 31.6 ±0.4; early blastocyst = 3.40± 67), compared to the control counterparts (morula = 6.2 ± 2.18; early blastocyst = 28.8 ± 2.69). This trend continued in subsequent days at E4.5, with most of the osmotically compressed embryos reaching late blastocyst stage at E5.5, suggesting that the delay did not compromise embryos viability. Next, to evaluate the impact of volumetric compression on embryo volume, we performed time-lapse imaging of PEG-treated embryos starting at the 4-cell stage. Embryos were stained with CellTracker, a nontoxic intracellular dye, and images using confocal three-dimensional (3D) scans at 30-minute intervals over a 5-hour period. PEG-treated embryos exhibited a sustained reduction in volume, indicating that volumetric compression persists over time rather than resolving quickly as would be expected in a purely osmotic shock (**Fig. 6c**). To further validate PEG effects on cytosolic mechanics, we conducted passive rheology measurements post-PEG treatment and found significant increase in cytosolic stiffness in both the 2-hour and 4-hour time points compared to controls (**Fig. 6d**, p < 0.01). This confirms that PEG treatment modulates cytoplasmic stiffness in line with volumetric compression.

**Figure 6.**
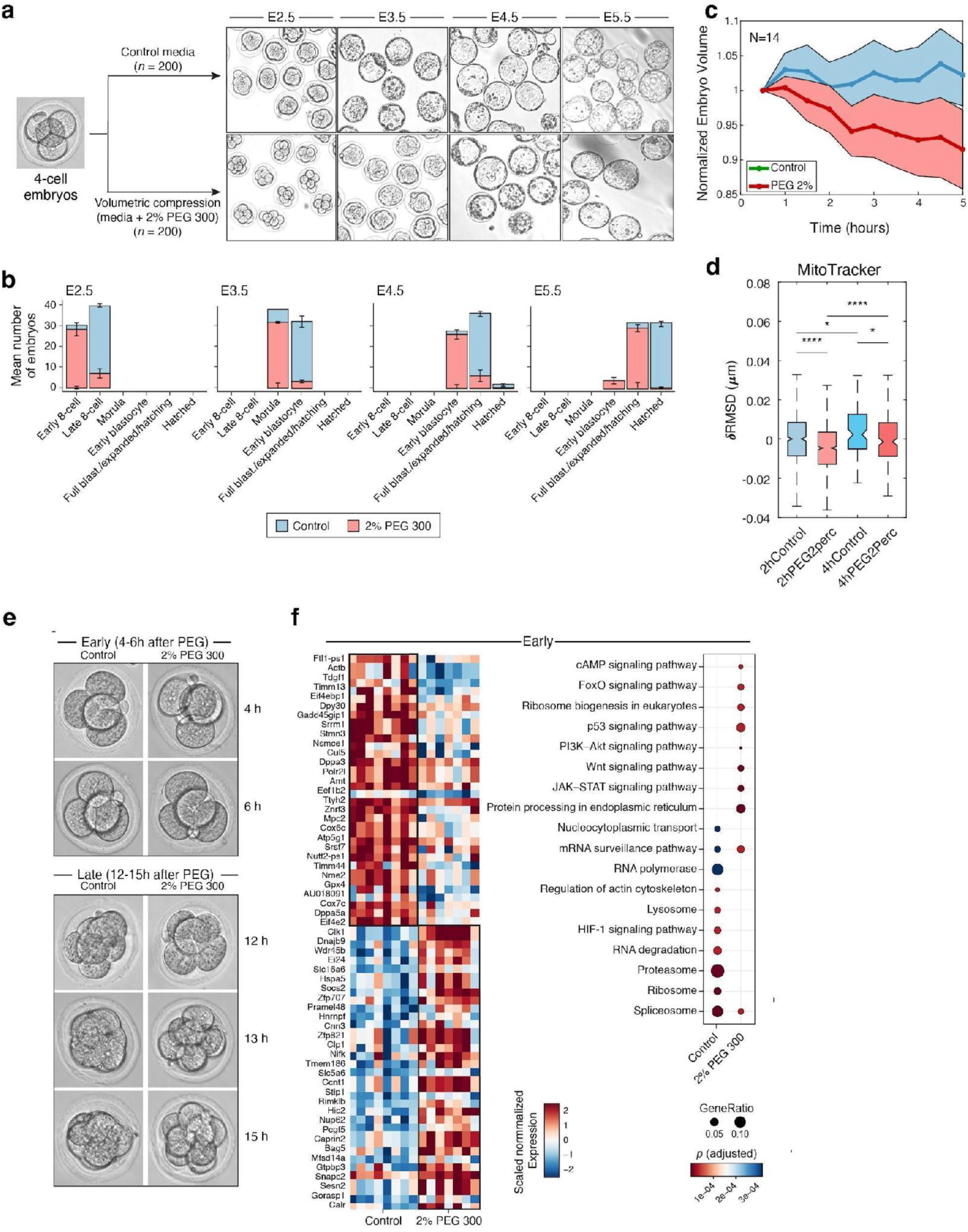
Volumetric compression causes delay in embryonic development. (**a**) Overview of mechanical perturbation (compression) experiment, with brightfield microscopy images of individual embryos. (**b**) Mechanical perturbation delays development. Number of embryos (y axis) observed at each stage (x axis) from control (blue) and compressed with 2% PEG (red) embryos. (**c**) Time-lapse imaging of compressed embryos starting at the early 4-cell stage. Embryos (control and volumetrically compressed, N=7 each) were stained with CellTracker and imaged over a 5-hour period. The volume of each embryo at each time point was normalized to its initial volume at the start of the experiment. Shaded regions represent 95% confidence intervals. (**d**) Passive rheology measurements using MitoTracker to assess mechanical properties of volumetrically compressed embryos. Distribution of δRMSD (y-axis) in control (blue) and volumetrically compressed (red) embryos at 2h and 4h post-treatment (x-axis). δRMSD, calculated as the deviation of RMSD from the 4-cell control reference stage, reflects relative differences in cytoplasmic softness. Boxplots denote medians and interquartile ranges (IQR), with whiskers representing the lowest and highest data points within 1.5 IQR of the lower and upper quartiles. Significance levels: ns (p > 0.05); * (p ≤ 0.05); ** (p ≤ 0.01); *** (p ≤ 0.001); **** (p ≤ 0.0001) based on the Wilcoxon ranked sum test. (**e,f**) Early expression changes associated with compression. (**e**) Representative phase-contrast images of control and 2% PEG 300-treated embryo groups (columns) harvested at specified time points (rows; n=4 per group and time point) for bulk RNA-seq of individual embryos. (**f**) Left: Expression (color bar, scaled normalized RNA level) of the top 30 genes (rows) differentially expressed between control and compressed embryos (columns) at the initial post-compression period (4, 6 h). Right: Significance (dot size, calculated using, Fisher’s exact test followed by p-value correction by Benjamini-Hochberg method) of enrichment of Gene Ontology terms (rows) among genes differentially induced in control or compressed embryos (columns).

To identify expression changes associated with the developmental delay induced by volumetric compression, we collected individual control and PEG300-treated embryos at 2-hour intervals starting from the 4-cell stage (**Fig. 6e**) and profiled individual embryos by plate-based full length whole-embryo RNA-seq (with SMART-seq2 ^63^). Because the developmental delay was evident by the 8-cell stage, we hypothesized that any causal expression changes precede it, possibly at 4-6 hours post-volumetric compression (while all embryos are in the 4-cell stage). Thus, we identified genes differentially expressed along the 4- to 8-cell developmental transition, by partitioning the control embryos into early (4-6h) and late (12-15h) time windows.

Comparing gene expression profiles between osmotically compressed and control 4-cell embryos at the 4-6 hour time window revealed 768 differentially expressed genes (**Fig. 6f**, adjusted p-value < 0.05; Bonferroni correction, **Supplementary Table 7**). Importantly, most (60%, 447 genes) of these differentially expressed genes were not among the 1,657 genes that are differentially expressed between early (4-6 hours, 4-cell stage embryos) and late (12-15 hours, 8-cell stage) control embryos (adjusted p-value < 0.05; Bonferroni correction, **Extended Data Fig. 13a,b Supplementary Table 6**). This suggests that the impact of compression on gene expression and embryo development is not merely a reflection of a delayed developmental trajectory (**Extended Data Fig. 13b**). Genes more highly expressed in control than compacted 4-cell embryos in the early window were enriched for proteasomal, lysosomal, autophagy, ribosome, and RNA degradation genes (**Fig. 6f**). Conversely, compression-induced genes were enriched for Hippo signaling, Wnt signaling, JAK-STAT signaling, mRNA surveillance, and AMPK pathway genes (not observed in genes differentially expressed between early and late control embryos), suggesting potential alterations in cellular signaling, growth, and metabolism induced by PEG300 treatment (**Fig. 6f**).

Interestingly, *Ctsl*, a lysosomal cysteine protease that is upregulated in early control 4-cell stage embryos and downregulated in control embryos later (**Extended Data Fig. 13c**), has lower expression in compressed early 4-cell embryos. This suggests that *Ctsl* inhibition can slow developmental progression, consistent with previous findings^64^. To directly test this, we inhibited lysosomal degradation by combining treatment with two two small molecules, E64d and pepstatin A (**Extended Data Fig. 13d**). E64d inhibits cysteine proteases such as cathepsins B, H, and particularly cathepsin L (Ctsl), while pepstatin A inhibits aspartic proteases like cathepsin D. We treated 20 embryos at the early 4-cell stage, alongside 20 controls. At E2.5, 70% of control embryos had reached the compacted stage and only 10% were in the early stage, whereas lysosome-inhibited embryos lagged significantly, with only 40% compacted, and 60% remaining in earlier stages. Notably, lysosome inhibition led to a similar developmental delay as seen in PEG-treated embryos, further suggesting that cytoplasmic de-crowding may be coupled to developmental progression.

Cytoplasmic crowding influences both the rate and equilibrium of molecular interactions ^59,61,62,65–67^, and we hypothesize that the increased spontaneous cytoplasmic motion we observed reflects a gradual reduction in molecular crowding throughout preimplantation development. A recent study in budding yeast described a phenomenon called “viscoadaptation”, in which cells actively stiffen their cytoplasm during temperature changes by converting glucose into storage carbohydrates (glycogen/trehalose) to enable homeostatic control of cellular viscosity to maintain constant rates of intracellular diffusion and diffusion-controlled chemical reactions^68^. In mice, total maternal glycogen declines from the oocyte to the early blastocyst and only partially rebounds when the trophectoderm establishes its own reserves, never reaching the oocyte’s high baseline ^69^. This parallels our finding of a gradual cytoplasmic softening from the 2-cell stage to blastocyst, with the trophectoderm consistently stiffer than the inner cell mass. Taken together, these observations suggest that tuning cytoplasmic viscosity might represent a conserved regulatory property—shaping processes as diverse as stress response in yeast and the timing of early embryonic development in animals.

## Discussion

Here, we developed a new methodology for simultaneous measurement of live mechanical properties, *in situ* gene expression in whole mount, and 3D morphology, and applied it to construct a multi-modal view of the earliest fate decisions in mouse embryos. This required two notable technical innovations: First, we developed scalable, multiplexed RNA *in situ* sequencing of thick 3D specimens along with high resolution antibody based imaging for segmentation and morphological feature extraction; and second, a novel approach for passive microrheology, leveraging the natural diffusional dynamics of endogenous organelles such as mitochondria and lysosomes, to noninvasively measure cytoplasmic stiffness in live embryos at single-cell resolution. Importantly, these organelles have readily available non-toxic live-cell imaging probes, allowing for highly scalable and facile measurements in 3D. These measurements were registered to spatial transcriptomic profiles within the same cells, resulting in the construction of a unified map of transcriptomic, morphological and molecular cell states (UTMM) within a single embryo. Applying UTMM to preimplantation embryo development, we extended previous works, by directly linking, for the first time, molecular, geometrical, and physical properties of cells during early lineage segregations.

Our current work provides the most extensive spatio-temporal single-cell profiling to date of preimplantation embryos covering the entire ICM-TE lineage segregation. While prior studies ^70^ that analyzed a limited number of single cells without spatial information identified ICM-TE lineage segregation in a subset of cells profile at the 16-cell stage, our data uncovers a gradual divergence of ICM and TE lineages as early as the early morula (<16 blastomeres), characterized by the expression of TE-specific (e.g. Cdx2) and ICM-specific (e.g. Sox2) gene programs. Notably, cells identified as ICM-like were generally closer to the embryonic center, while TE-like cells were predominantly peripheral. However, a considerable overlap in the radial spatial distribution of ICM-like and TE-like cells and the ICM and TE scores at this stage suggests that transcriptional changes emerge gradually, with spatial bias becoming more pronounced over time, and both expression states and positioning strengthening concurrently.

The origin and timing of the initial differentiation of the Inner Cell Mass (ICM) into epiblast (Epi) and primitive endoderm (PrE) lineages remain under active investigation ^31^. Previous single-cell studies reported limited or inconsistent separation of presumptive Epi and PrE cells within early inner-cell populations, with robust segregation emerging only at later blastocyst stages^71^. In particular, Ohnishi et al. analyzed a small number of E3.25 ICM cells (33 cells from four embryos, assayed for ∼50 genes) and did not observe reproducible clustering into PrE- and Epi-like groups, whereas E3.5 ICM cells did segregate, consistent with progressive lineage resolution. Using multiplex *in situ* measurements of ∼150 genes across hundreds of pre-cavitation inner cells (278 cells from 17–32-cell embryos), combined with NMF-based program discovery that captures continuous mixtures of transcriptional states rather than requiring discrete clusters, we detect reproducible early lineage-biased transcriptional programs within morula inner cells. The larger gene panel, increased cell numbers, and program-level analysis increase sensitivity to subtle coordinated transcriptional shifts that may precede overt lineage segregation. We therefore interpret ICM-1 (“Epiblast-biased”) and ICM-2 (“PrE-biased”) not as committed lineages, but as early transcriptional biases that precede the discrete Epi and PrE segregation observed at later blastocyst stages. In our data, ICM-1 cells predominantly occupy internal positions, with 85% exhibiting minimal exposed surface area, indicating a strong predisposition for internalization and stabilization within the ICM. In contrast, ICM-2 cells, with 35% showing higher surface exposure (>0.25), indicating that a sizable fraction of ICM-2 cells are observed while still partially exposed at the embryo surface and likely undergoing inward repositioning/internalization ^30,31,40,46^. This transcriptional bias is detectable before morphogenetic events such as blastocyst cavity formation, consistent with recent live-imaging lineage tracing showing the emergence of an early SOX2⁺ epiblast-biased population at the late morula stage together with continued lineage switching at later stages, highlighting early lineage bias in the presence of persistent plasticity ^72^. Earlier blastocyst-stage work also reported mosaic Nanog/Gata6 expression and early lineage bias within the E3.5 ICM prior to overt Epi/PrE segregation ^33^. Together, these results show that transcriptional priming of ICM-derived lineages, particularly the ICM-2 program, can arise in cells that are not yet fully internalized, indicating a partial decoupling between spatial position and transcriptional state. We therefore interpret these data as evidence that positional rearrangements and lineage-biased gene expression programs emerge in overlapping temporal windows, rather than following a strict hierarchical sequence. ^33^

For the TE lineage, TE-1 and TE-2 cells exhibit transcriptional biases consistent with early polar- and mural-like programs already detectable at the mid-late morula stage. Both subtypes remain spatially peripheral, with similar radial distances from the embryo center; however, after aggregation and computational alignment of embryos into a common coordinate system, we observe reproducible spatial polarization, with TE-1 and TE-2 cells enriched on opposite sides of the embryo. Importantly, these patterns reflect probabilistic transcriptional biases rather than fixed lineage commitment, as extensive plasticity is known to persist through these stages. We therefore interpret this polarization as an early organizational tendency that precedes cavitation, indicating that molecular priming and spatial organization can emerge in parallel before lumen formation.

Our work establishes a novel framework for measuring cytoplasmic mechanics, enabling single-cell, whole-mount, non-invasive assessments that span both external and internal cells throughout embryogenesis. While earlier studies ^7,39,40^ have elegantly detailed cortical mechanics, including softening during cleavage stages up to early 8-cell and tension increases upon 8-cell compaction, their focus on cortical properties and external cells provides a complementary perspective. Our findings reveal a continuous cytoplasmic softening from the 4-cell to morula stage, with no significant difference between early and compacted 8-cell embryos, suggesting that cytoplasmic viscoelasticity provides distinct and biologically relevant insights into intracellular mechanics, which directly influence molecular crowding and mobility. Importantly, the early lineage segregation of morula cells in the 9-16 cell stage coincides with differential mechanical properties aligning with their distinct developmental programs, with ICM-like cells softer than TE-like cells. At the mid-late morula stage, ICM-2 cells were stiffer (lower RMSD values) than ICM-1 cells, in agreement with recent evidence from optical-tweezers experiments, showing that the cytoplasm of PrE-primed mESCs are stiffer than that of EPI-primed mESCs^73^. We emphasize that detecting early transcriptional or mechanical biases does not imply deterministic fate specification. Classical experiments demonstrate substantial lineage plasticity in early embryos, and our data are fully consistent with this view. Rather than predicting final fate, UTMM reveals subtle, probabilistic biases that emerge alongside morphogenetic events and may influence, but do not determine, later lineage outcomes.

We also demonstrated the generality of our method by deploying our organelle-tracking passive microrheology approach to measure cytoplasmic stiffness of NSCs in adult acute brain sections, followed by post facto immunofluorescence to measure NSC state. Our finding that activated NSCs (aNSCs) exhibit greater lysosomal displacement than quiescent NSCs (qNSCs), indicative of a softer cytoplasm, is consistent with an emerging link between intracellular mechanical state and NSC activation. In a complementary study, Bressan et al. demonstrated that ependymal cilia-driven mechanical signals maintain NSC quiescence through PKD1/2–TRPM3-mediated Ca²⁺ transients, and that CRISPR-mediated deletion of Trpm3 in qNSCs reduces Ca²⁺ entry and triggers their transition toward an activated, proliferative state^74^. Strikingly, applying the same organelle-tracking framework to Trpm3-knockdown tissue, showed a reproducible shift in organelle MSD toward greater compliance, consistent with cytoplasmic softening upon reduced Ca²⁺-driven cytoskeletal contractility. Together, these results suggest that cytoplasmic stiffness is not merely a passive correlate of NSC activation state, but may reflect the downstream mechanical output of mechanotransduction pathways that regulate the quiescence–activation balance.

While organelle-based passive microrheology provides a non-invasive and endogenous method for measuring cytoplasmic mechanical properties, it has certain limitations that merit consideration. First, organelles exhibit active, motor-protein-driven transport in addition to thermal fluctuations, particularly over longer timescales. To overcome this, our analysis is restricted to high-frequency, short time scale fluctuations, where thermal motion dominates. However, the presence of active transport mechanisms could still introduce variability that requires careful data interpretation. Second, the size, shape, or distribution of organelles may vary across different developmental stages and cell types in different tissues, potentially influencing measurements. This may especially be the case for mitochondria, which are subject to fission, fusion and interactions, and whose degree of oblateness varies significantly across developmental stages and tissues ^75^. During preimplantation development, mitochondria are structurally immature, characterized by a predominantly spherical morphology and sparse cristae ^55,76,77^. As differentiation progresses, mitochondria transition to elongated forms with developed cristae, forming filamentous and reticular networks, reflecting shifts in metabolic state and bioenergetic demand of the cells ^75^. In our work, we demonstrate that mechanical measurements from mitochondria, lysosomes, and beads are highly correlated, suggesting an approach to bypass organelle specific measurement biases. However, further validation and benchmarking are needed to apply this approach for other tissues and developmental stages. Third, passive microrheology cannot directly capture cortical tension or mechanical forces at the membrane, which are equally critical for understanding the overall role of mechanics in cell fate, techniques like micropipette aspiration remain necessary complements. Lastly, the reliance on fluorescent staining introduces potential photobleaching and phototoxicity concerns, particularly for extended live-imaging experiments, which could limit the temporal resolution or experimental duration. Addressing these limitations through benchmarking with complementary techniques will further enhance the robustness and applicability of organelle-based passive microrheology for developmental studies.

We observed that volumetric compression delayed developmental progression in 4-cell stage embryos. Cytoplasmic crowding can affect the rate and equilibrium of molecular interactions ^59,61,62,65–67^. We hypothesize that the increase in spontaneous cytoplasmic motion we observed may indicate a continuous decrease in molecular crowding along preimplantation embryogenesis. We further hypothesize that the mechanical and physical environment of the cytoplasm may be a ‘regulatory clock’ that helps coordinate the pace and timing of the developmental progression. In this model, signaling pathways may depend on molecular transport, which is sensitive to molecular de-crowding. Since volumetric compression is expected to lead to crowding and prevent softening, this may further support this model. In our study, we utilized PEG osmolarity compression, a standard approach in the field ^49,60,67,78^, to reduce cellular volume, however, osmolarity compression also introduces transient ionic imbalances which may also contribute to developmental delay. The reduction in expression of lysosomal genes in compressed 4-cell embryos may point at a role for lysosomal and related pathways, including ubiquitin-mediated degradation and autophagy, in de-crowding. Notably, lysosomal degradation of maternal proteins is employed extensively from the 2-cell to the morula stage, and interference with this pathway is lethal at the 4- to 8-cell stage ^79,80^.

In conclusion, our work represents a new class of joint measurements of molecular and physical properties, including mechanics and geometry, assessed spatially. Such approaches will be critical for understanding and engineering 3D biological tissues. Recent studies with 3D *in vitro* models, including organoids and *in vitro* grown synthetic embryos, have sought to recapitulate developmental processes, but have large systematic variability, and, often, variable efficiencies and reproducibility. Multi-modal measurements such as UTMM may help us understand and control variation in these systems to enable engineering of more complex and accurate *in vitro* models. Beyond development, the coupling of mechanics, geometry, and gene expression may be especially important in understanding tissue pathophysiology in tumorigenesis, fibrosis and wound healing. Future advancements of the UTMM technology in scalability, measurement modalities, and throughput will enable us to generate physical-molecular models to learn the rules of tissue organization in health and disease.

## Acknowledgments

This work was supported by the National Institutes of Health (grant nos. R01HG010647 to F.C.). F.C. also acknowledges support from the Searle Scholars Award, the Burroughs Wellcome Fund CASI award, and the Merkin Institute. A.S. was supported by the NIH Neuroimaging Training Program T32 grant 5T32EB001680. L.L. was supported by the National Science Foundation Graduate Research Fellowship under Grant No. 2141064. ESB was supported by NIH 1R01EB024261, HHMI, Lisa Yang, John Doerr, NIH R01MH124606, and NIH 1U19MH114821.

## Author contributions

In this study, conceptualization was led by EH, MG, AR, and FC, who together defined the research goals and strategy. EH, AS and HY performed all the experiments. EH, AS, PY, HY, FC, LL, and ZDC conducted the analysis. DS, ESB, MG, AR, and FC provided essential resources. Code development and adaptation were carried out by EH, AS, PY, HY, ZDC, LL, and DAW. EH, AS, PY, and HY validated the results for accuracy and reproducibility. The methodology, including statistical and analytical techniques, was developed by EH, AS, PY, HY, and YL. EH, AS, and HY were responsible for the investigation process. Visualization of the data was managed by EH, AS, PY, and HY, with EH, AS, PY, and HY also taking charge of data curation. Funding acquisition was secured by MG, AR, FC, and ESB, with MG, AR, and FC providing project supervision. The writing of the original draft was primarily undertaken by EH, AS, HY, MG, AR, and FC, while all authors contributed to reviewing and editing the manuscript, ensuring its scholarly integrity and compliance with publication standards.

## Data and materials availability

Processed is deposited for review: https://drive.google.com/drive/folders/1P6k7sq5Ltzl9ZfEwgILcJGy1uDJN0m8l?usp=drive_link. Processed gene expression matrices will be deposited to single-cell portal. Sequence data will be deposited on SRA. Code for data processing and to reproduce analyses are available on github https://github.com/ehabibi1/UTMM/tree/main. Raw data will be available online after publication.

## EXTENDED DATA FIGURE LEGENDS

**Extended Data Figure 1.**
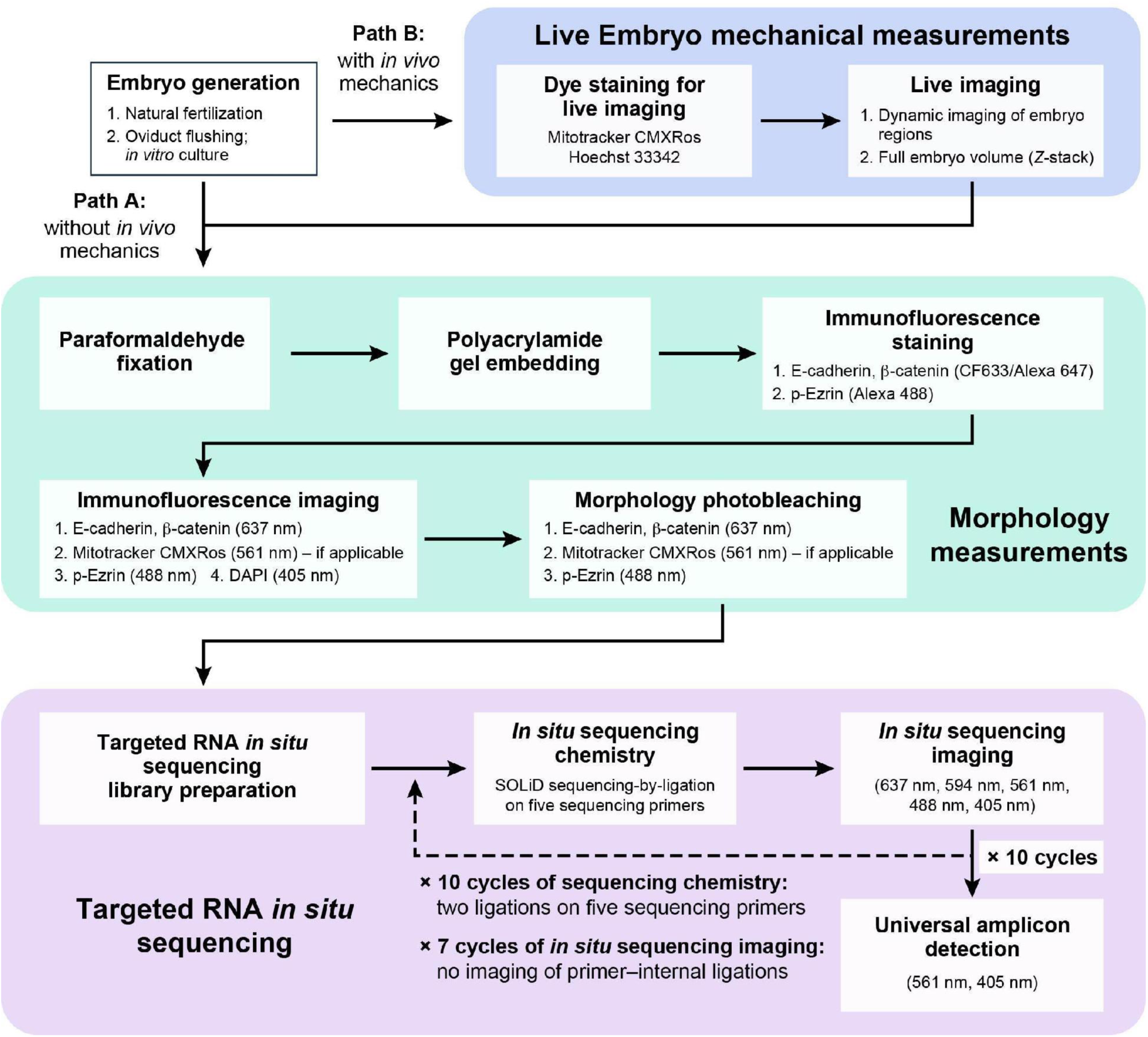
Experimental and analysis workflow. The initial workflow for embryos that were not mechanically characterized is in Path A. UTMM workflow is in Path B.

**Extended Data Figure 2.**
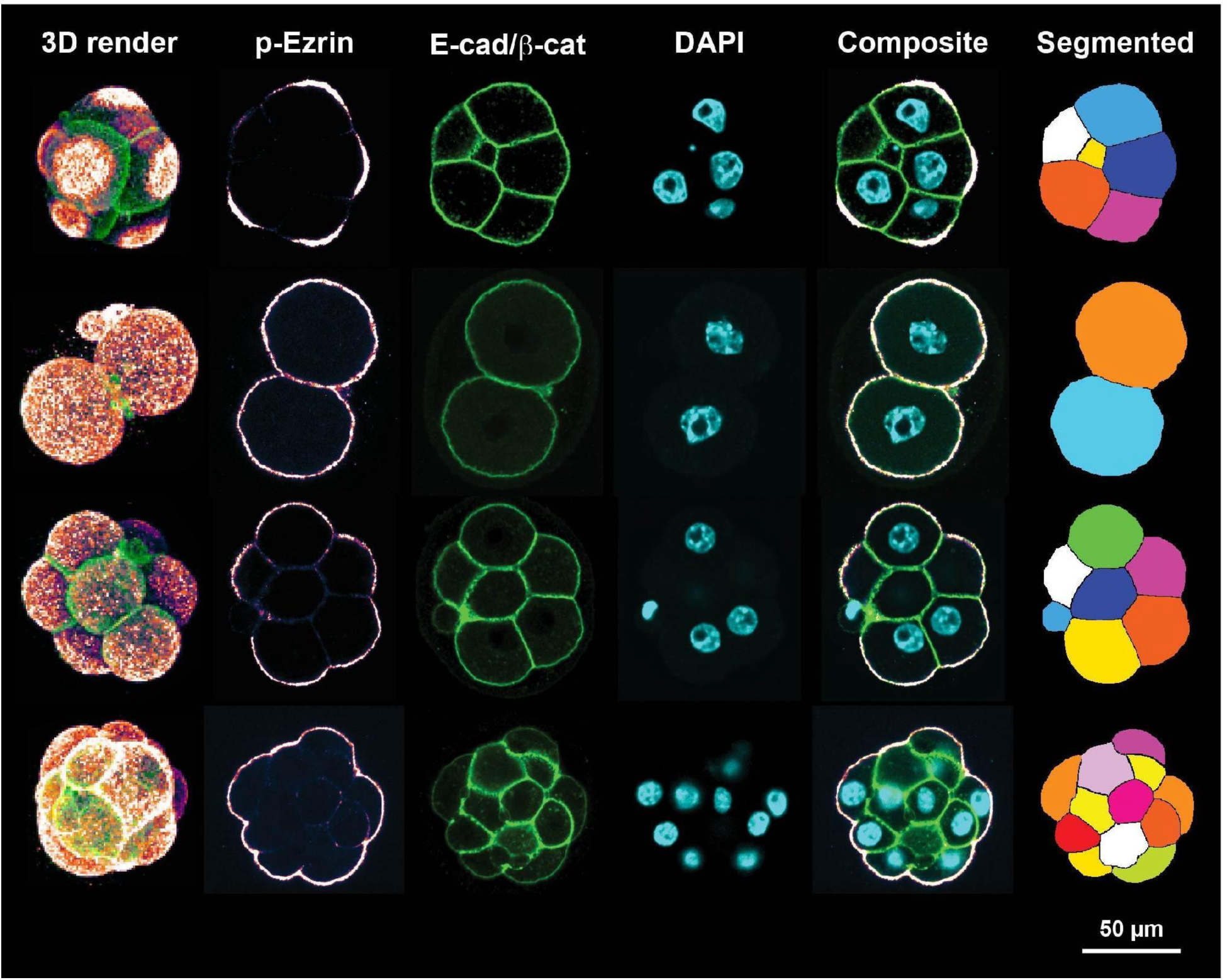
Immunostaining and 3D segmentation. From left: Confocal microscopy images of immunostaining and computed cell segmentation masks for embryos at various stages immunostained for p-Ezrin, E-cadherin and β-catenin in the same color channel (green), and DAPI (cyan). Leftmost: maximum intensity projections of confocal z-stacks; all other columns: a single z-plane. Rightmost: segmented masks, pseudocoloring cells to highlight morphology. Scale bar: 50 μm.

**Extended Data Figure 3.**
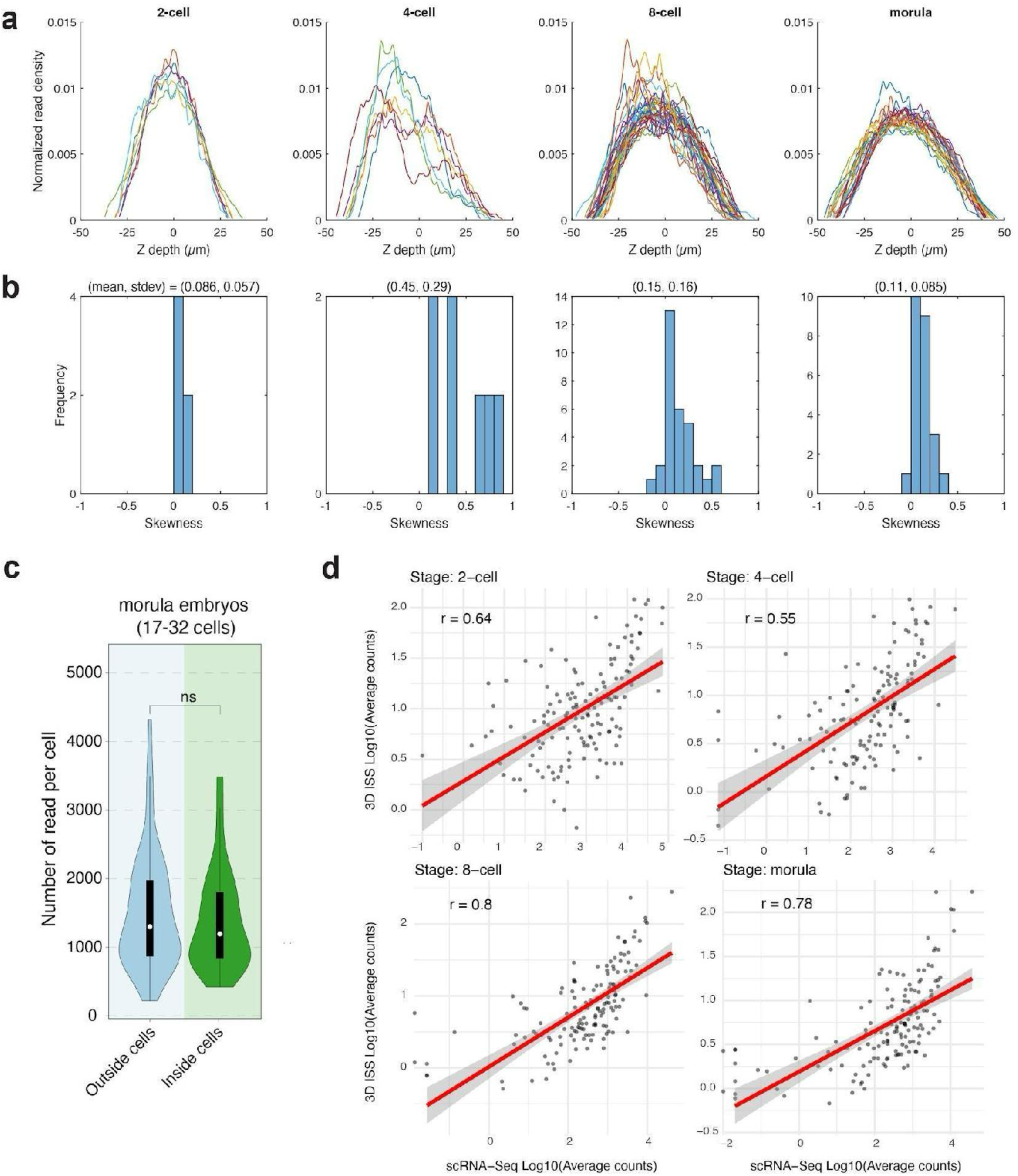
Quality of 3D volumetric in situ sequencing. (**a**) Normalized read densities (y axis) at different z-section depth (x axis) for each embryo in the dataset (colored line) in each stage (columns). (**b**) Distribution of skewness (x axis) of read z-profiles across stages (columns, **Methods**). Higher skewness values and variation at the 4-cell stage is attributable to the asymmetry and orientation of 4-cell embryos. As the embryo becomes more sphere-shaped through the morula stage, the skewness decreases. (**c**) Comparison of read counts between internal and external cells in 3D *in situ* sequencing. Internal and external cells were defined based on the Euclidean distance from each cell to the center of the embryo (**Methods**). The violin plot displays the distribution of detected read counts, demonstrating comparable efficiency of transcript detection across both internal and external cell populations within the embryo. ns, p > 0.05 ; *, p ≤ 0.05 ; **, p ≤ 0.01 ; ***, p ≤ 0.001, ****, p ≤ 0.0001, Wilcoxon ranked sum test. (**d**) Correlation of averaged gene expression estimates between 3D *in situ* sequencing (3D ISS) and single-cell RNA sequencing at matched stages (log10-transformed counts on both axes, r represents Pearson’s correlation coefficient).

**Extended Data Figure 4.**
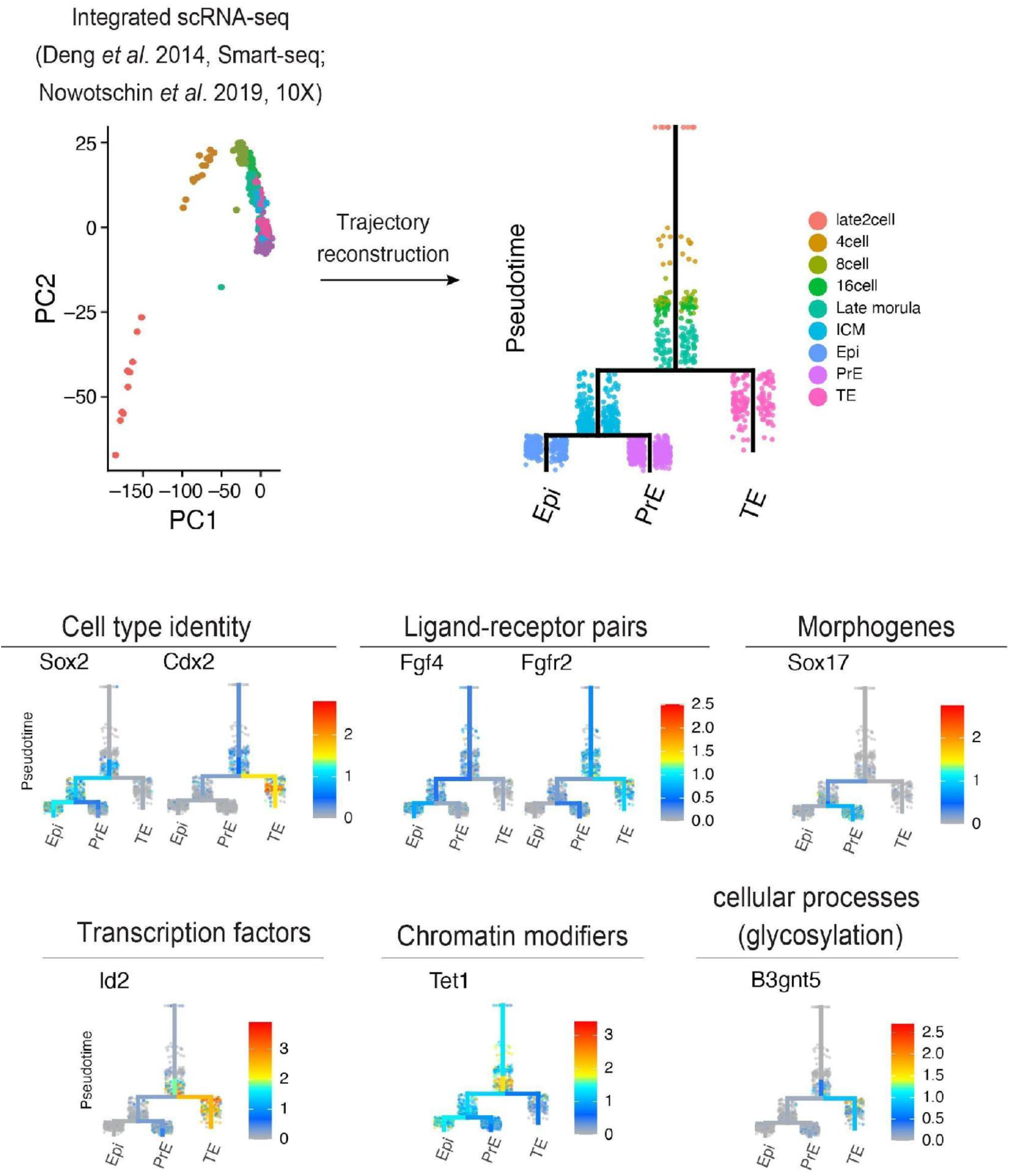
Trajectory reconstruction of cell profiles along mouse preimplantation development. Top left: scRNA-seq profiles (dots) embedded in the space of the first two principal components (PCs) of a PCA, colored by developmental stage. Top right and bottom: The same profiles (dots) along a branching tree reconstruction by URD^25^, colored by developmental stage (top right) or by expression (colorbar) of key genes (bottom).

**Extended Data Figure 5.**
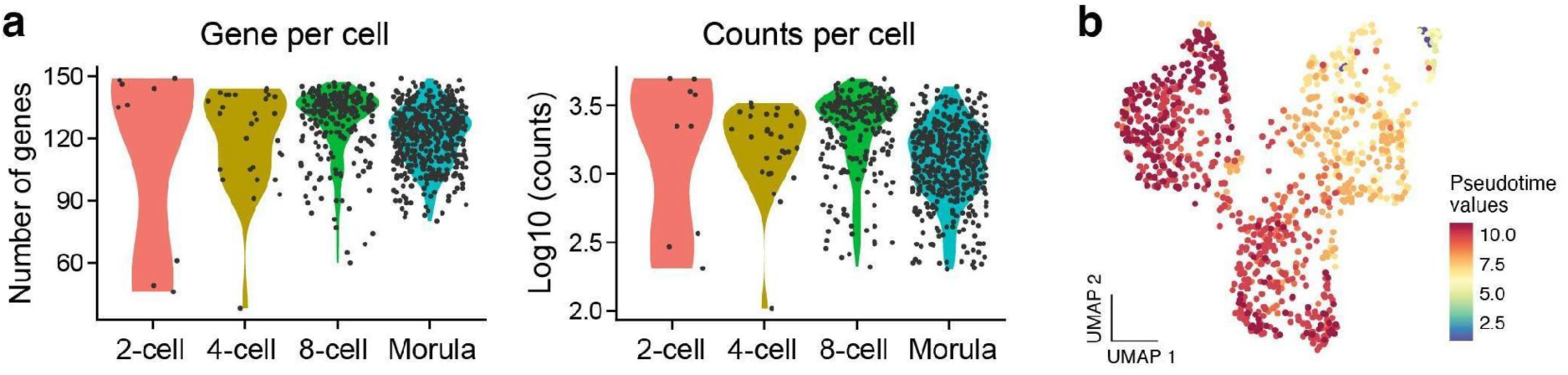
Quality assessment of *in situ* cell profiles. **(a)** Distribution of number of genes and mRNA molecules (y axis) called per cell (dot) by targeted in situ sequencing across developmental stages (x axis). **(b)** UMAP embedding of *in situ* RNA cell profiles (dots) colored by a diffusion pseudotime approach (color bar, **Methods**).

**Extended Data Figure 6.**
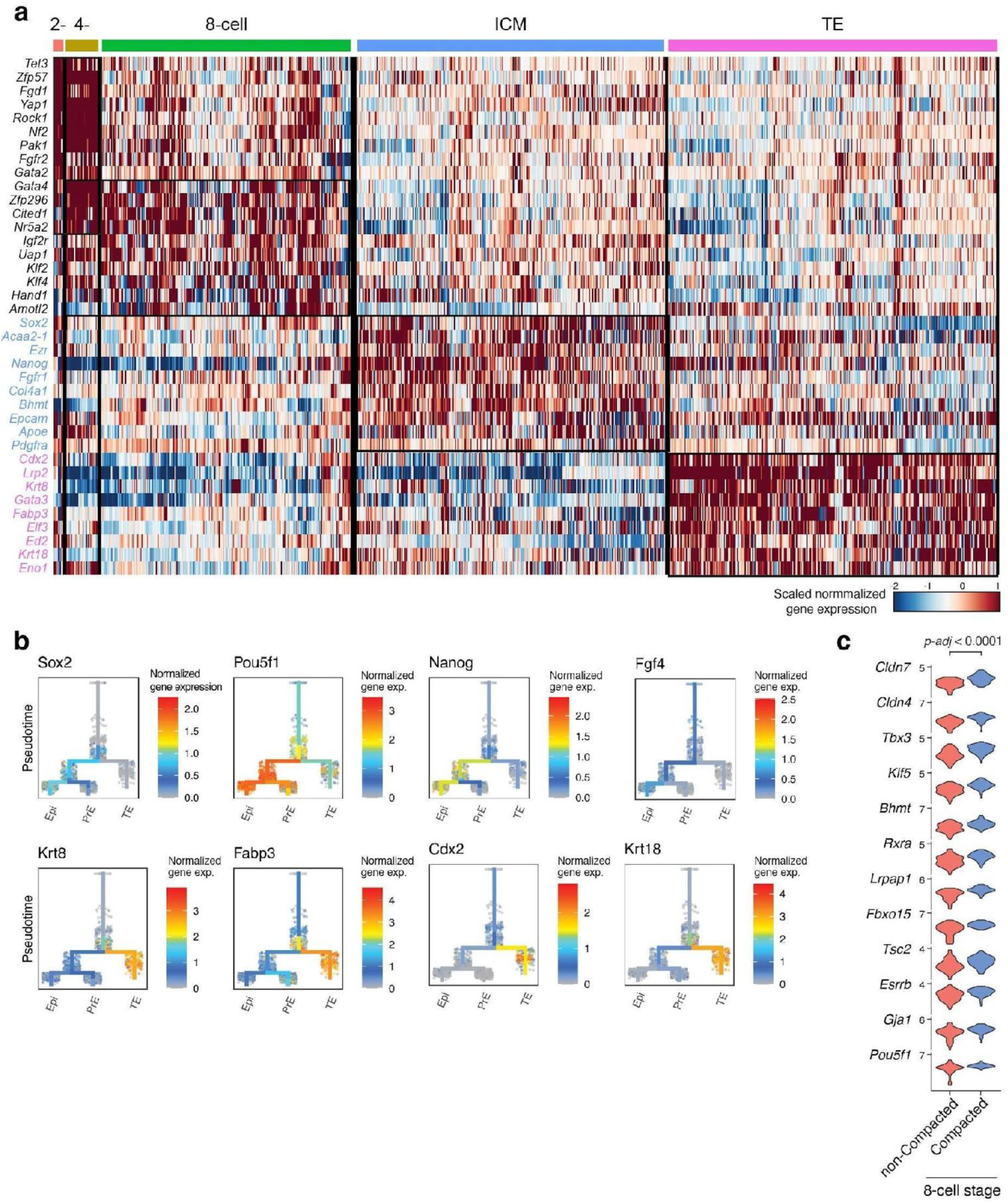
Distinct expression signatures associated with cell states in preimplantation embryos. (**a**) Relative expression (scaled normalized gene expression) of top 10 differentially expressed genes (rows) between *in situ* cell profiles (columns) of each stage (color code on top). (**b**) scRNA-seq profiles (dots) along a branching tree reconstruction by URD^25^(as in **Extended Data** Fig. 4), colored by expression of select ICM (top) and TE (bottom) markers. (**c**) Distribution of expression (y axis, normalized gene expression) of top differentially expressed genes between cells from compacted and non-compacted 8-cell stage embryos (x axis).

**Extended Data Figure 7.**
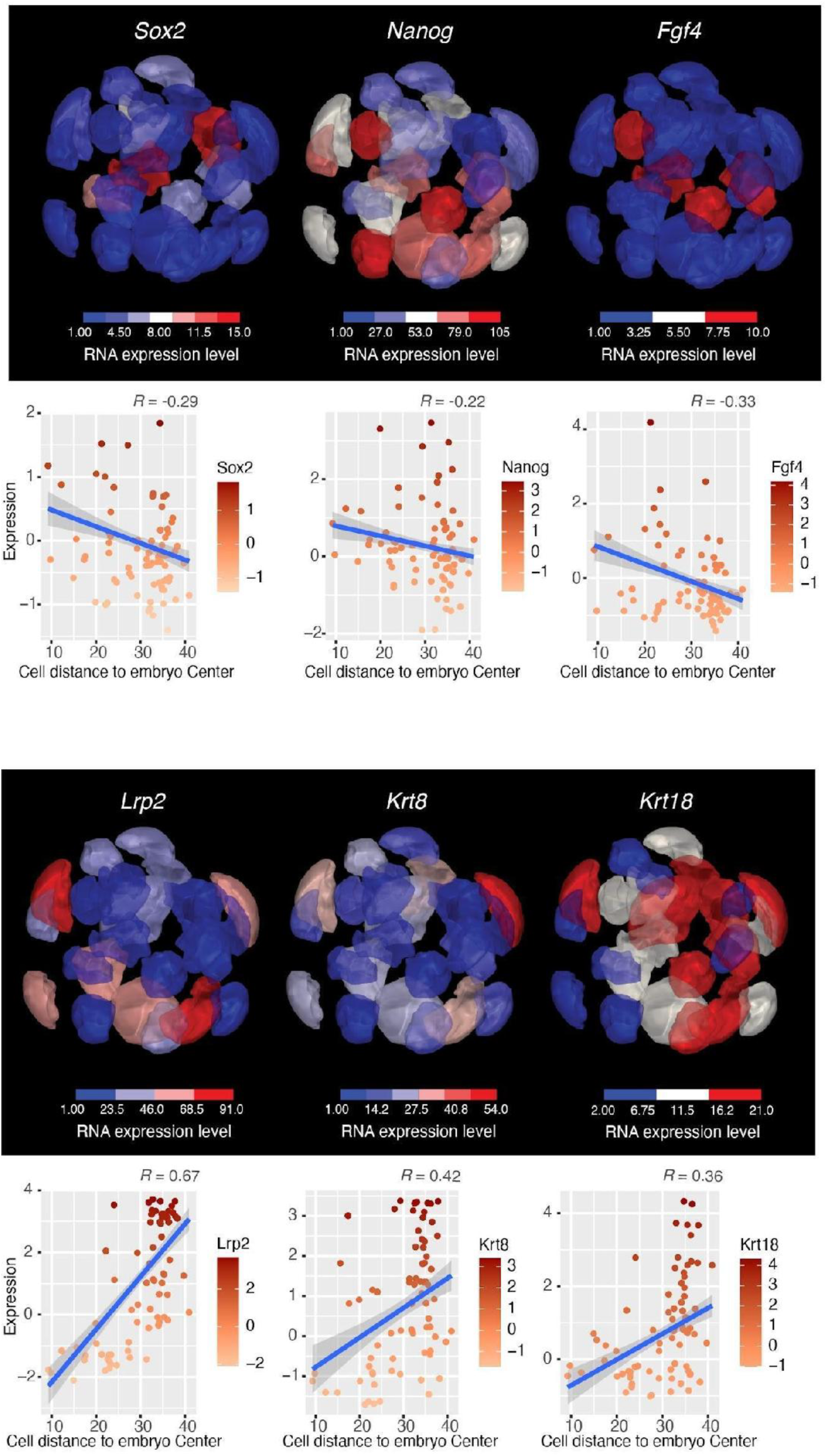
Genes expression associated with cell position in 16-cell stage embryo. Top: Expression level (transcript count per cell) of canonical markers in each cell of one 16-cell stage embryo (enhanced visualization by separating the individual cells). Bottom: Expression (x axis, scaled normalized gene expression) of the same select genes and distance to embryo center (y axis, Euclidean distance) in each cell of 16 cell stage embryos.

**Extended Data Figure 8.**
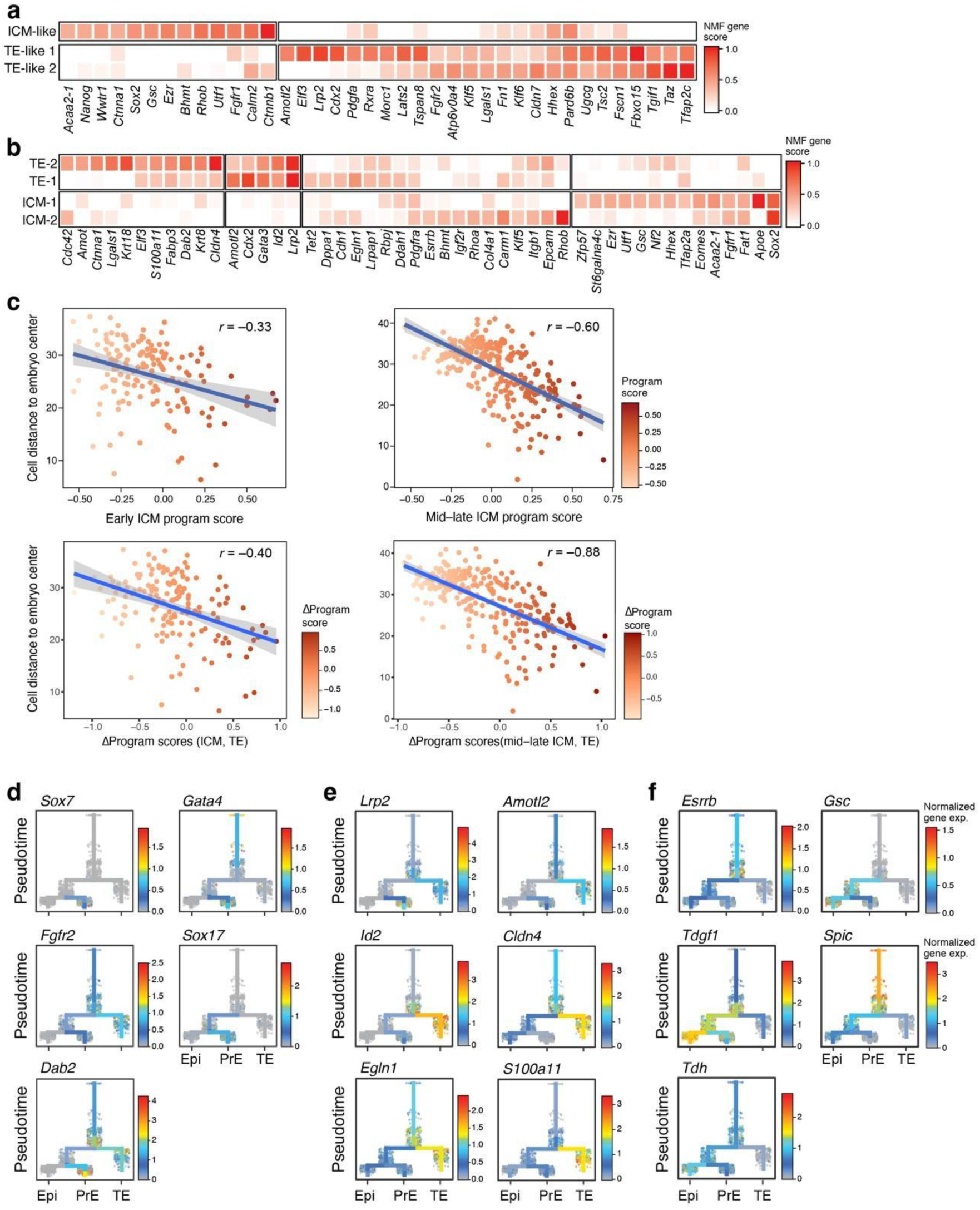
Spatial distribution of TE and ICM cell subsets. (a,b) NMF program. NMF gene weights (color bar) of top genes (columns) for each program (row) defined at early (a) or mid (b) morula. (**c**) ICM program scores (top, x axis, and dot color) or difference between ICM and TE scores (bottom, x axis and dot color) and distance from embryo center (y axis) for each cell (dot) in early (left) or mid (right) morula embryos. (**d-f**) scRNA-seq profiles (dots) along a branching tree reconstruction by URD^25^(as in **Extended Data** Fig. 4), colored by expression of select PrE (**d**), TE (**e**), and Epi (**f**).

**Extended Data Figure 9.**
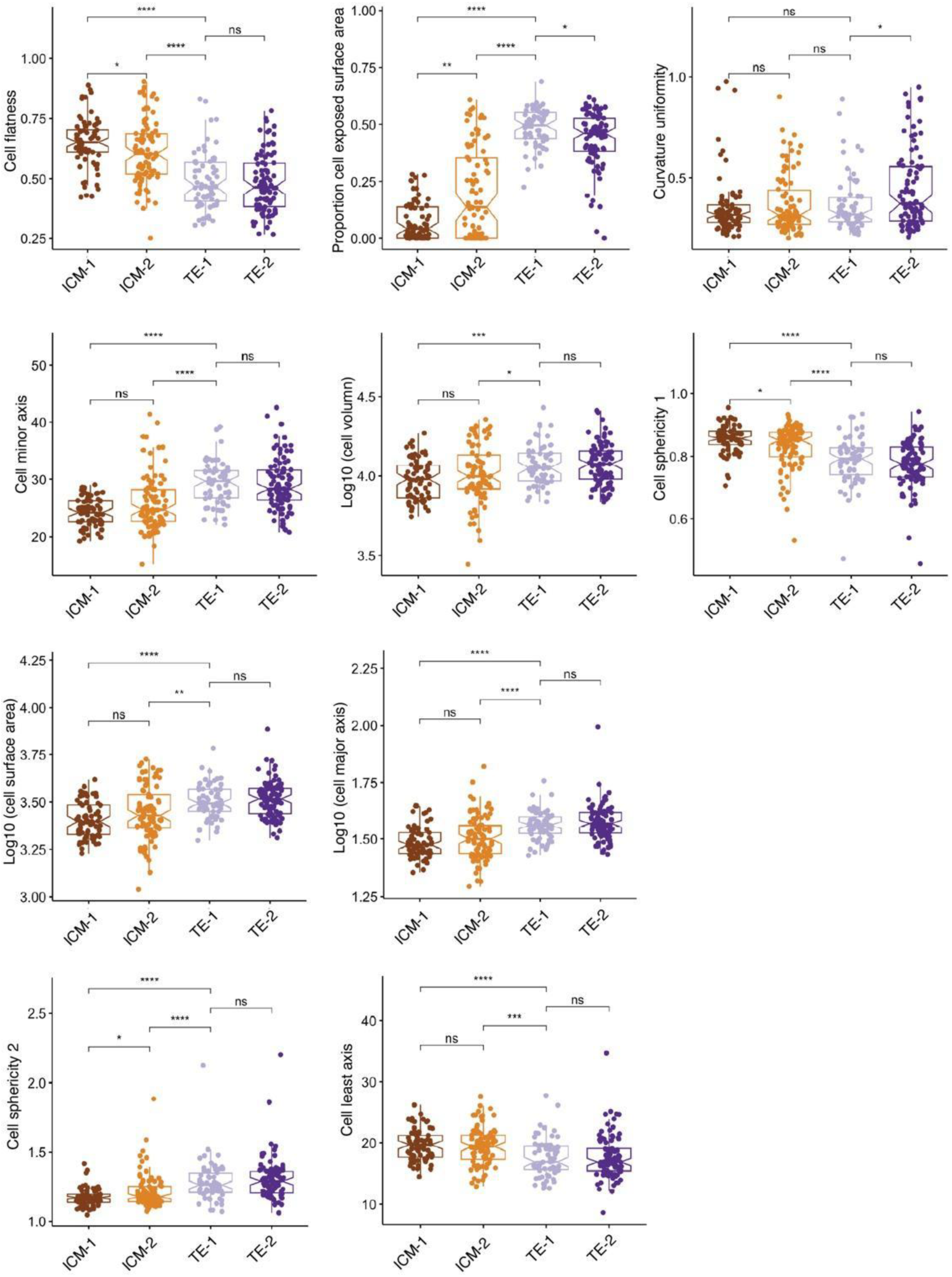
Expression cell states are associated with different morphological characteristics. Distribution of different morphological features (y axis) in each of four mid-morula expression cell states (x axis). ns, p > 0.05 ; *, p ≤ 0.05 ; **, p ≤ 0.01 ; ***, p ≤ 0.001, ****, p ≤ 0.0001, Wilcoxon ranked sum test.

**Extended Data Figure 10.**
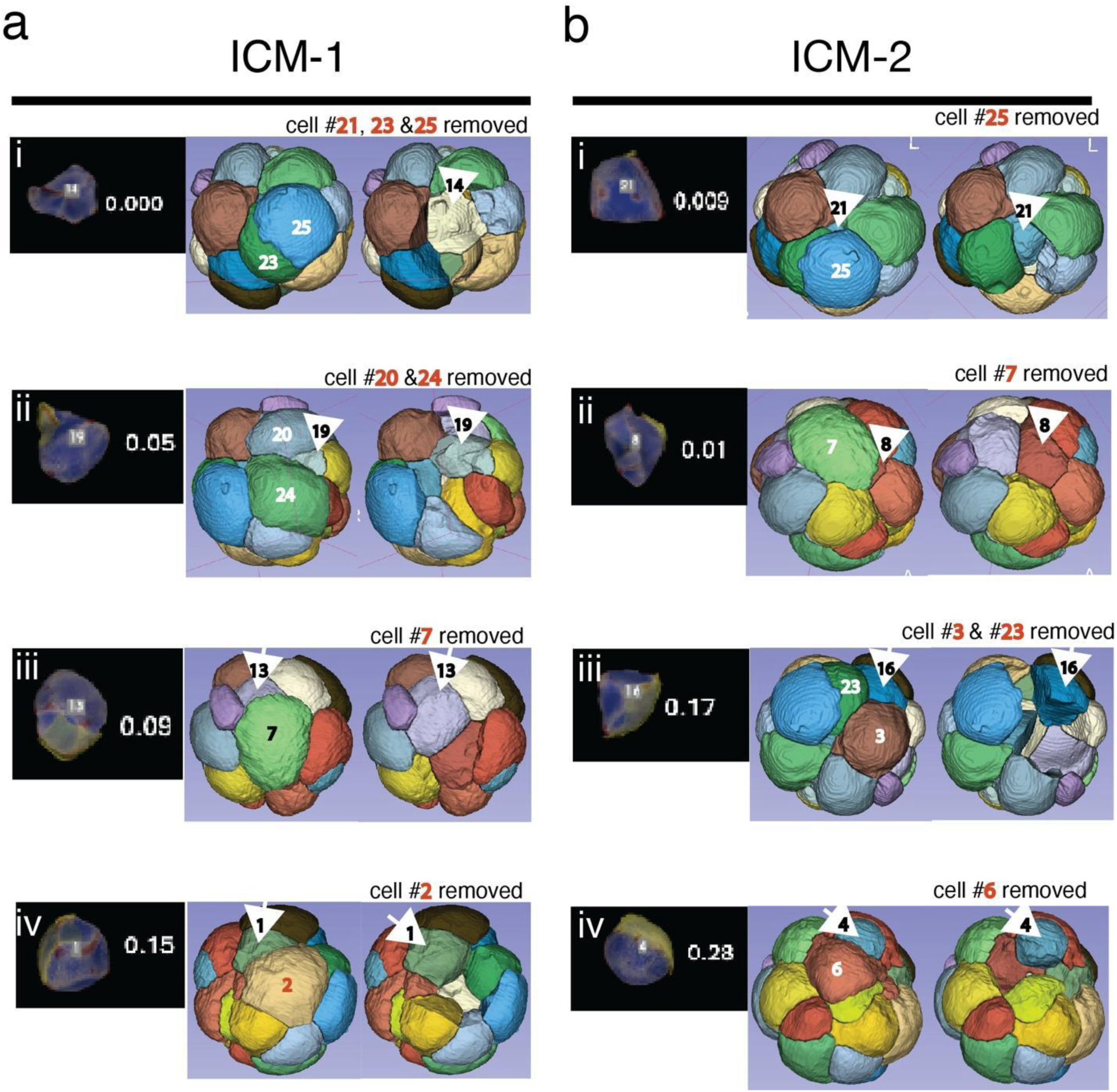
Spatial and morphological variability of ICM-1 and ICM-2 cells during early ICM development. Representative examples of (**a**) ICM-1 and (**b**) ICM-2 cells across a range of exposed surface areas. Individual cells are shown against a black background, color-coded by scores for unexposed (blue) and externally-exposed (yellow) cell surface area, and curvature (red), with the calculated surface area exposed to the outside of the embryo displayed numerically. (i-iv) For each column, panels show the progression of exposed surface areas from minimal exposure (i) to higher exposure values (iv). To the right, corresponding 3D reconstructions of the morula embryo depict the spatial location of the selected cells. In the rightmost column, for clarity, neighboring cells have been partially removed in the 3D view to emphasize the positioning of the target cells within the embryo.

**Extended Data Figure 11.**
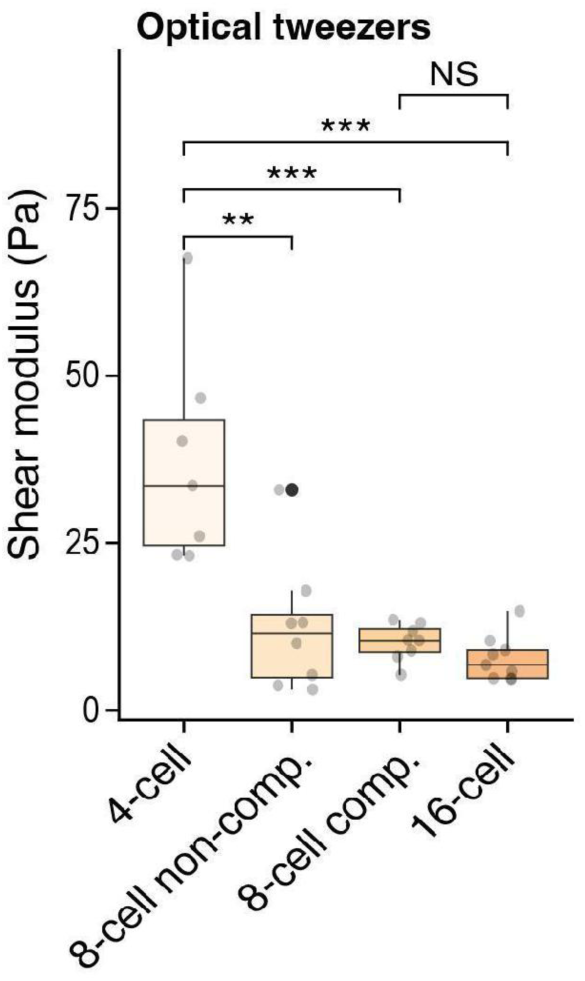
Microrheology validation using endocytosed microbeads. Distribution of shear modulus measured by optical tweezers (y axis) in each development stage (x axis).

**Extended Data Figure 12.**
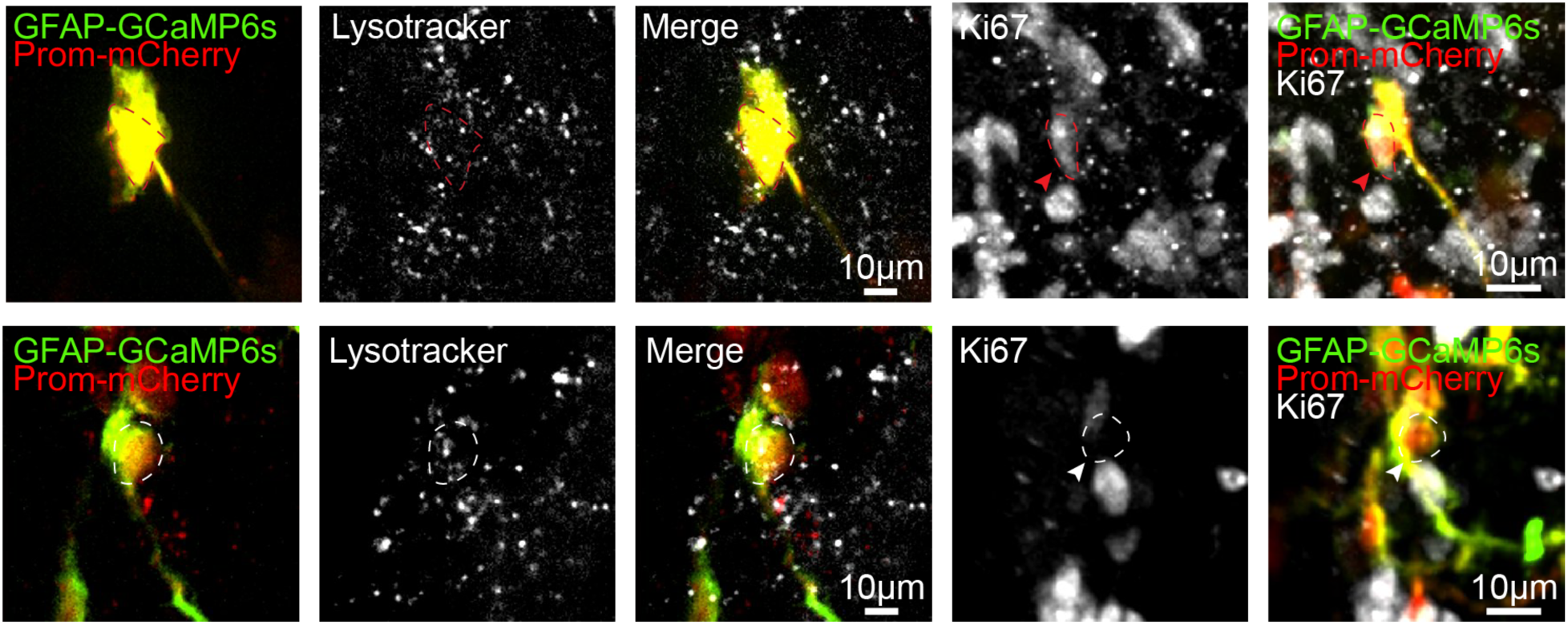
Representative images of activated and quiescent neural stem cells in the postnatal mouse brain. Representative images of lysosomes (white) in activated neural stem cells (aNSCs) and quiescent neural stem cells (qNSCs) imaged in acute brain slices at 74.6 frames/s. Post hoc identification of imaged NSCs was performed by Ki67 immunolabeling. Red arrowheads and ROIs indicate Ki67+ aNSCs, whereas white arrowheads and ROIs indicate Ki67− qNSCs.

**Extended Data Figure 13.**
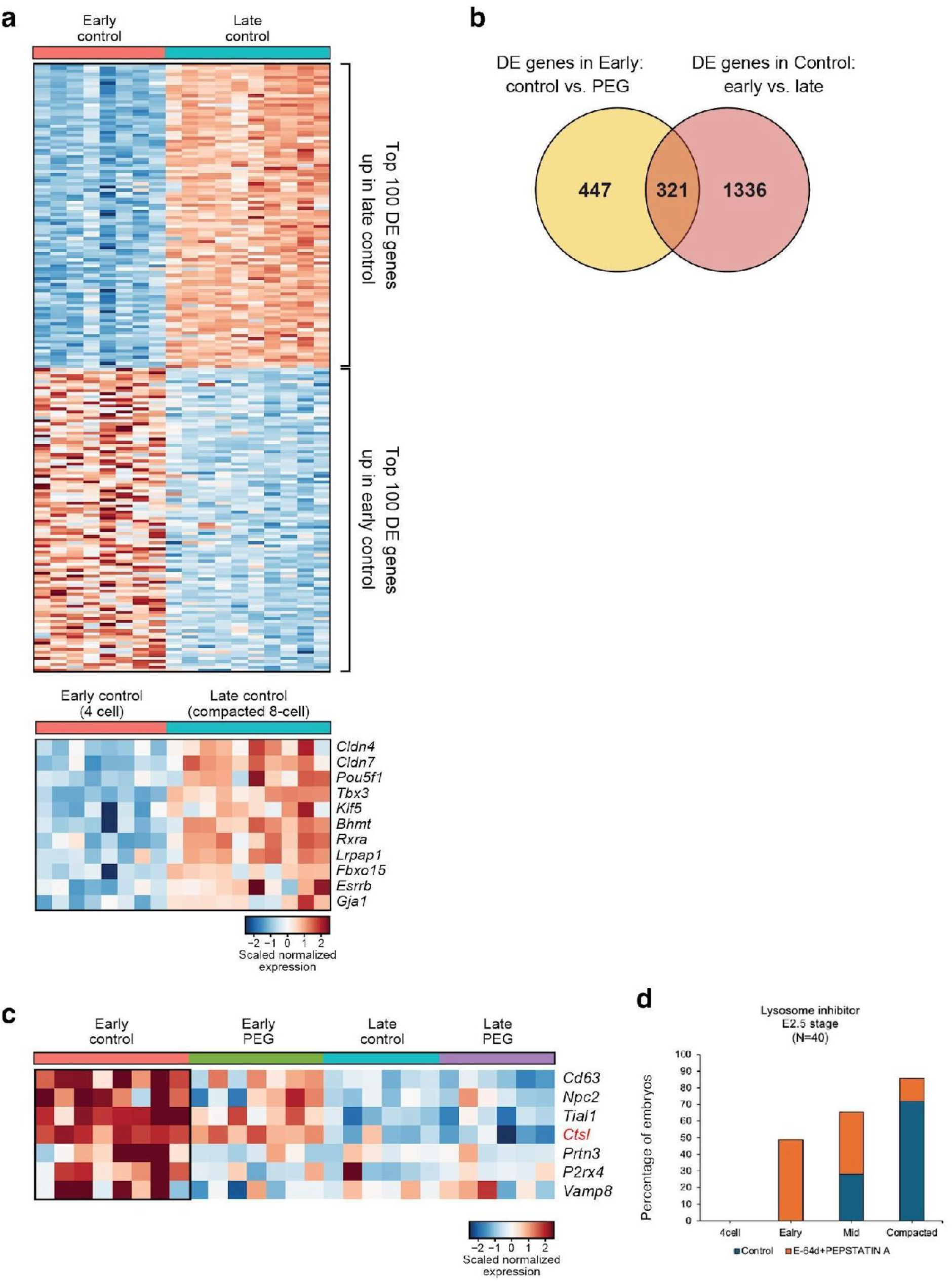
Gene expression analysis in compressed embryos. **(a)** Top: Expression (scaled normalized gene expression) of the top 100 differentially expressed genes between early and late developmental windows in each of the early or late control embryos in SMART-seq data (columns, **Supplementary Table 8** for list of genes). Bottom: Expression (scaled normalized gene expression) of each of the genes (rows) upregulated at late 8-cell stage as identified by RNA in *situ* sequencing data as shown previously in Extended Data Figure 6c **(b)** Overlap between genes differentially expressed in early PEG-treated *<u>vs</u>*. early control embryos (both directions) and in early vs. late control embryos (both directions). **(c)** Expression (scaled normalized gene expression) of genes (rows) associated with the lysosomal compartment in control and PEG-treated groups embryos (columns) at early and late stages (color label bar). (**d**) Inhibition of lysosomal degradation delays development. Percentage of embryos (y-axis) in each developmental stage by E2.5 (x-axis) in control (blue) and E64d+Pepstatin A treated (orange) groups (where treatment commenced at the 4 cell stage).

## Live-cell mechanics measurements

Embryos with mechanical measurements were characterized as follows.

### Staining

For each cohort of live-cell measurements, a fresh stock of 1 mM MitoTracker Red CMXRos (ThermoFisher, M7512) in DMSO was prepared, following the manufacturer’s instructions. Embryos were then stained with 100 nM MitoTracker Red CMXRos and 5µg/ml Hoechst 33342 (Biotium 40046) in M2 or KSOM media for 60-120 minutes. For experiments with LysoTracker Deep Red, embryos were stained with 100 nM LysoTracker Deep Red as above. For experiments with tracer particles, the zona pellucida was first dissolved by treating 2-cell stage embryos with X for Y minutes, followed by staining with 0.5-μm-diameter carboxylate-modified polystyrene latex beads (1:2000 dilution; Sigma, L3280). After staining, embryos were briefly washed with M2 media, then transferred to M2 media under mineral oil in a 35 mm glass-bottom Petri plate (Cellvis, D35-14-1.5-N) for imaging.

### Imaging

Dynamic imaging of live embryos (64 x 64 pixels, 23 x 23 μm, imaged every 12 ms for 8 seconds) was performed using a Leica SP8 point-scanning confocal microscope with a resonant scanner. A 63X/NA 1.20 water immersion objective was used. Color channels were as follows: 405 nm at 2% intensity (DAPI) and 552 nm at 1% intensity (MitoTracker) lasers used to excite the sample, and HyD detectors used for detection (410-547 nm for DAPI, and 561-763 nm for MitoTracker). Measurements were acquired from multiple locations within each embryo. Stage positions were recorded for each of the measurements for data registration.

Following dynamic imaging, a volumetric Z-stack of the entire embryo was acquired (1024 x 1024 pixels, 148 x 148 μm in XY, a voxel size of 0.244 μm from the bottom to top of the embryo). The same 63X/NA 1.20 objective was used. Color channels were as above, except with increased laser power (405 nm at 20% intensity for DAPI and 552 nm at 5% intensity for MitoTracker). Embryos were then fixed, as described above.

## Immunofluorescence and *in situ* sequencing

Briefly, fixed embryos were embedded in a polyacrylamide gel to keep them immobilized during subsequent immunofluorescence staining and imaging, and targeted *in situ* sequencing library preparation and sequencing. After gelation, the terms “embryos” and “gels” are used interchangeably in the experimental steps. All reagents used through the end of library preparation were RNAse-free. All volumes for solutions added to the sample gel are 300 μL, except for inexpensive buffer-only solutions (i.e. 1X PBS, 2X SSC, wash buffer, etc.), which are used at a volume of 1000 uL.

### Embedding embryos into polyacrylamide gel

The gel-embedding approach was adappTEd from *in situ* genome sequencing (IGS) ^22^ and implemented as described below.

Wells of 24-well glass-bottom plates (Cellvis, P24-1.5H-N) were treated with Bind-Silane (GE Healthcare, GE17-1330-01) as previously described ^21^, with the following modifications: a 1:250 dilution of Bind-Silane was used, and the reaction time with Bind-Silane was extended to 45 minutes. To prepare the lids of the gelling chambers by making them hydrophobic, cut pieces of glass microscope slides (approx. 8 mm x 8 mm) were treated with Sigmacote (Sigma SL2) for 1 minute, then washed three times with water. To form spacers for gelation, four layers of Invisible Tape (Universal, 83412, total spacer thickness = 4 x ∼60 um = ∼240 um) were attached to the cut glass pieces near two opposite edges on the same face of the glass, with a gap between the spacers ^22^. Combined lid/spacer units were prepared weeks in advance, and set aside until use.

Gelling monomer mix was prepared on ice by mixing 40 μL of 40% 19:1 acrylamide:bis-acrylamide (ThermoFisher, AM9022), 100 μL of 4X PBST (4X PBS (ThermoFisher, AM9625), 0.4% Tween-20 (Teknova, T0710, 10% Tween-20 stock)), and 244 μL H_2_O (ThermoFisher, 10977015), forming 384 μL of monomer mix. To cast the gel, 8 μL of 10% TEMED (Sigma T7024, 0.2% final concentration) and 8 μL of 10% APS (ThermoFisher, 17874, 0.2% final concentration) were added to the monomer mix in that order, vortexing in between each addition, forming the final gelling mix, which was kept on ice and used immediately.

To cast the gel, two droplets of gelling mix were pipetted into the lid of a 35 mm Petri dish, and 10 μL of gelling mix were added to the center of the well of the 24-well glass bottom plate (treated as described above). Embryos were washed with gelling mix by transferring them from storage buffer after fixation (1X PBS with 0.1% PVP), to the first droplet, then transferring them to the second droplet, before transferring them to the droplet of gelling mix in the plate.

To track embryos while minimizing the total number of samples, embryos that were characterized in the live state were grouped together into sets that contained only one embryo of a given developmental stage. Each set was washed together, and added to the same droplet region in the plate. Through careful handling, multiple sets were embedded into the same gel, with > ∼1 mm separation between sets. Embryo handling was performed under a stereo microscope.

After embryo transfer, gelling chamber lids were lowered with forceps into the well on top of the droplet. Plates were placed on a heating block set to 45°C for 2 minutes to initiate gelation and minimize motion. The plate was then placed into a humidified plastic container, which was purged with nitrogen gas for 10 minutes at room temperature. The container was sealed, and incubated for 2 hours at 37°C. After gelation, the lid glass was removed with forceps, and the gel was washed with 1X PBS for 10 minutes at room temperature. The gel was washed again with 1X PBS, and stored at 4°C.

### Embryo permeabilization

Gel-embedded embryos were permeabilized with 1% Triton X-100 in 1X PBS for 30 minutes at room temperature, then washed twice with 1X PBS at room temperature for 15 minutes each wash.

### Immunofluorescence – staining

Embryos were blocked by washing with 1% (w/v) UltraPure BSA (ThermoFisher, AM2618) in 1X PBS for 2 hours at 4°C. Immunostaining for E-cadherin (ThermoFisher, 13-1900 [host: rat], 1:200 dilution; and Clone 36, BD Biosciences, 610181 [host: mouse], 1:200 dilution), Beta-Catenin (clone 15B8, Abcam ab6301, [host: mouse], 1:200 dilution) and p-Ezrin (Cell Signaling Technology 3141S, [host: rabbit], 1:400 dilution) was performed in 1% BSA in 1X PBS overnight at 4°C. On the next day, embryos were washed 5 times for 30 minutes each wash with 1X PBS at room temperature. Immunostaining with secondary antibodies (Donkey anti-Rat CF633 [MilliporeSigma, SAB4600133]; Goat anti-Mouse Alexa 647 [ThermoFisher, A21235]; Goat anti-Rabbit Alexa 488 [ThermoFisher, A11034]; all antibodies at 1:500 dilution) was then performed in 1% BSA in 1X PBS overnight at 4°C. Signals for E-cadherin and Beta-catenin were merged into the 647 color channel. Following secondary antibody staining, embryos were washed 5 times for 30 minutes each wash with 1X PBS at room temperature. Embryos were then stained with 2 mg/L DAPI (ThermoFisher, D21490) in 1X PBS for 1 hour at room temperature, then washed 2 times for 30 minutes each wash with 1X PBS.

### Immunofluorescence – imaging

Morphology was imaged on an Andor Dragonfly Spinning Disk confocal microscope (described below) using the following color channels, with associated power and exposure times: 637 nm (30% power, 300 ms), 488 nm (30% power, 300 ms), and 405 (100% power, 100 ms). For embryos that were characterized with MitoTracker CMXRos, residual dye was also imaged in the 561 nm channel (100% power, 100 ms). Imaging was performed with 0.4 μm Z-steps.

### Immunofluorescence – bleaching

Following imaging, the morphology signal in the 637 nm and 488 nm color channels was photobleached by re-imaging the sample in laser widefield mode using the following color channels, with associated power and exposure times: 637 nm (100% power, 4000 ms), 488 nm (100% power, 3000 ms). For embryos that were characterized with MitoTracker CMXRos, residual dye was also photobleached by imaging in the 561 nm laser widefield channel (100% power, 8000 ms).

After photobleaching, embryos were imaged again (as described above for imaging immunofluorescence in confocal mode) to confirm photobleaching. Then, embryos were washed with 1X PBS for 15 minutes at room temperature, followed by a brief digestion with 0.2 U/mL Proteinase K (NEB) in 1X PBS for 10 minutes at room temperature. Samples were then washed 3 times for 15 minutes each wash with 1X PBS at room temperature.

### 3D targeted *in situ* sequencing library preparation

Library preparation was adapted from Expansion Sequencing ^21,82^, with modifications as noted below. Embryos were pre-hybridized with wash buffer (20% formamide (ThermoFisher, AM9344) in 2X SSC (20X SSC stock, ThermoFisher 15557044)) for 30 minutes at room temperature, then incubated for 24 hours with hybridization mix. The hybridization mix consisted of 105.5 μM pooled probe set (total of 2,344 padlock probes, ∼45 nM each probe; normalized from probe set subpools 1-3 described above) in 20% formamide, 2X SSC buffer.

Following hybridization, samples were washed five times for 30 minutes each wash with wash buffer at 37°C, followed by one wash for 30 minutes with 1X PBS at 37°C. Embryos were then washed with 1X SplintR Ligase Buffer (NEB, M0375L) for 30 minutes at room temperature. Pre-ligation was performed by incubating the embryos with ligation mix (1250 U/mL SplintR Ligase (NEB, M0375L) in 1X SplintR Ligase Buffer) for 6 hours at 4°C. Samples were then ligated by incubating the embryos with freshly prepared ligation mix overnight at 37°C. After ligation, samples were washed with 2X SSC for 30 minutes at room temperature, followed by a pre-hybridization wash with wash buffer for 30 minutes at room temperature. Samples were then incubated with 500 nM rolling circle amplification primer (RCA primer, **Table S3**) in wash buffer for 3 hours at 37°C. Samples were then washed with wash buffer for 30 minutes at 37°C, followed by 1X PBS for 30 minutes at 37°C.

Rolling circle amplification was then performed. Samples were pre-equilibrated with 1X EquiPhi29 DNA Polymerase Reaction Buffer with 1 mM DTT (ThermoFisher, A39392) for 30 min at room temperature. Pre-amplification was performed by incubating embryos with pre-amplification mix (1000 U/mL EquiPhi29 DNA Polymerase, 1 mM DTT in 1X EquiPhi29 DNA Polymerase Reaction Buffer) for 6 hours at 4°C. Rolling circle amplification was then performed by incubating embryos with amplification mix (1000 U/mL EquiPhi29 DNA Polymerase, 250 μM each dNTP (NEB, N0447L), 40 μM amino-allyl dUTP (ThermoFisher, R1101), 25 nM compaction oligo (**Table S3**, see ^84^ for details), and 1 mM DTT in 1X EquiPhi29 DNA Polymerase Reaction Buffer) for 16 hours at 30°C. After amplification, samples were washed for 30 minutes with 1X PBS at room temperature, then treated with 5 mM BS(PEG)9 (ThermoFisher, 21582) in 1X PBS for 2 hours at room temperature. Samples were then washed with 1 M Tris pH 8.0 (ThermoFisher, AM9855G) for 15 minutes at room temperature, then washed twice with 1X PBS for 15 minutes each wash at room temperature. Embryos were then washed with strip buffer (80% formamide (ThermoFisher, 4311320) in ddH_2_O) 3 times for 10 minutes each wash at room temperature to strip the compaction oligo, followed by 4 washes with 1X PBS for 5 minute each wash at room temperature. Rolling circle amplification products are called amplicons or RCA colonies (rolonies).

### *In situ* sequencing – preparation and overview

The chemistry and approach for *in situ* sequencing was modified from ExSeq and IGS ^21,22^. The formulation of cleavage buffers was adoppTEd from OligoFISSEQ ^85^.

Embryos were washed with open instrument buffer (150 mM Tris base (Sigma, T6066), 100 mM sodium acetate (Sigma, S2889), 0.05% Tween 20, pH to 7.4 with glacial acetic acid (VWR, 9508-03)) for 4 times for 5 minutes each wash at room temperature. To dephosphorylate 5’ phosphates, samples were then treated with Quick CIP solution (250 U/mL Quick CIP in 1X rCutSmart Buffer (NEB, M0525L)) for 45 minutes at room temperature. Samples were then washed with an open instrument buffer for 5 times for 5 minutes each wash at room temperature.

*In situ* sequencing using SOLiD Sequencing-by-Ligation was then performed. A total of 10 ligations, two consecutive ligations on each of the five sequencing primers (SeqN-1, SeqN, SeqN-2, SeqN-3, SeqN-4, **Table S3**) were performed. The first ligations were imaged only on SeqN-1 and SeqN, while the second ligations were always imaged. Thus, a total of 7 rounds of sequencing imaging data were collected.

For each sequencing primer, the following steps were performed: primer hybridization; first ligation; DAPI staining and imaging first ligation (only for sequencing primers SeqN-1 and SeqN); dephosphorylation; cleavage; second ligation; DAPI staining and imaging second ligation (all primers); and stripping.

While this protocol used SOLiD sequencing oligos, which are no longer commercially available, equivalent oligonucleotides can be purchased from IDT, as previously detailed ^22,85^.

### *In situ* sequencing – chemistry

Sequencing primers were hybridized by incubating the sample with 2.5 μM sequencing primer in 5X SASC (0.75 M sodium acetate (Sigma, S2889), 75 mM sodium citrate (Sigma, S1804), pH 7.5) for 45 minutes at room temperature. Samples were then washed 3 times for 5 minutes each wash with an open instrument buffer at room temperature.

Ligation was then performed. The sample was treated with ligation mix (60 U/μL T4 DNA Ligase (Rapid) [Enzymatics, L6030-W-L], 1 mg/mL UltraPure BSA, 1:400 diluted SOLiD sequencing oligos (ThermoFisher, 4475669) in 1X T4 DNA Ligase Buffer) for 90 minutes at room temperature, then washed 3 times for 10 minutes each wash with open instrument buffer.

For sequencing primers SeqN-1 and SeqN, DAPI staining and imaging of the first ligation was performed (see below). For other sequencing primers (SeqN-2, SeqN-3, and SeqN-4), samples proceeded directly to dephosphorylation.

Dephosphorylation was then performed as described above (treated with Quick CIP solution, 250 U/mL Quick CIP in 1X rCutSmart Buffer for 45 minutes at room temperature, followed by 5 washes for 5 minutes each wash at room temperature).

SOLiD oligos cleavage was then performed, using open formulations of SOLiD sequencing buffers ^85^. The sample was washed 2 times for 5 minutes each wash with cleavage buffer C (50 mM AgNO3 (Sigma, S8157) in H_2_O) at room temperature, followed by 2 washes for 5 minutes each wash with cleavage buffer B (50 mM 2-mercaptoethanolsulfate (Sigma, M1511), 30 mM trisodium citrate, 300 mM sodium acetate, pH 7.5) at room temperature. The sample was then washed 3 times for 5 minutes each wash with open instrument buffer. Second ligation was then performed, with the same ligation conditions as the first ligation (above).

After the second ligation, imaging was performed (below). Then, sequencing products were stripped by washing with strip buffer (80% formamide in water) 4 times for 10 minutes each wash at room temperature, followed by washing with open instrument buffer 3 times for 5 minutes each wash, before proceeding to the next primer hybridization.

### *In situ* sequencing – imaging

Each imaging cycle began by staining the sample with 2 mg/L DAPI in an open instrument buffer for 1 hour at room temperature. The sample was then washed with an open instrument buffer for 30 minutes before imaging. GLOX imaging buffer was then prepared immediately before use ^22^.

Briefly, ∼200 μL of GLOX master mix was prepared (56 mg/mL glucose oxidase (Sigma, G2133), 1:5 diluted catalase (Sigma, C3155) in dilution buffer (200 mM Tris pH 8, 50 mM NaCl)). GLOX imaging buffer was then prepared from 20% (w/v) glucose, dilution buffer, 200 mM Trolox (Sigma, 238813-5G) stock (in anhydrous DMSO, ThermoFisher, D12345) and GLOX master mix, with a final composition of: 1:50 diluted GLOX master mix in 10% glucose, 94 mM Tris pH 8.0, 23.5 mM NaCl, and 2 mM Trolox. 2 mL of GLOX imaging buffer was prepared and added to the well, and replaced with fresh GLOX imaging buffer after every ∼60-90 minutes of imaging.

Imaging was performed on an Andor Dragonfly Spinning Disk confocal microscope (below) using the following color channels, with associated power and exposure times for imaging the first ligation (on SeqN-1 and SeqN sequencing primers): 637 nm (100% power, 100 ms), 594 nm (100% power, 100 ms), 561 nm (100% power, 300 ms), 488 nm (50% power, 400 ms), and 405 nm (100% power, 100 ms). For imaging the second ligation (on all sequencing primers), the following laser power and exposure times were used: 637 nm (100% power, 150 ms), 594 nm (100% power, 150 ms), 561 nm (100% power, 450 ms), 488 (50% power, 450 ms), and 405 nm (100% power, 100 ms). Imaging step size was 0.4 μm.

After imaging each ligation, the sample was washed 4 times for 5 minutes each wash at room temperature with an open instrument buffer, before proceeding to dephosphorylation and cleavage (post-first ligation), or stripping (post-second ligation).

### Universal rolony (amplicon) detection hybridization

After *in situ* sequencing, including stripping of the final *in situ* sequencing products, a universal rolony detection probe (**Table S3**) that binds to the constant region of all amplicons in the specimen was hybridized at 100 nM for 1 hour in 1X PBS. The sample was then washed 3 times with 1X PBS for 15 minutes each wash. Imaging buffer was then prepared (as described above), added to the sample, and imaged on the Andor Dragonfly Spinning Disk confocal microscope with the following color channels and powers/exposure times: 561 nm (100% power, 200 ms), and 405 nm (100% power, 100 ms). Imaging step size was 0.4 μm. This dataset is referred to as the rolony detection hybridization round.

### Microscope configuration

For imaging embryo morphology and *in situ* sequencing, an Andor Dragonfly spinning disk confocal (CR-DFLY-201-40) coupled to a Nikon Ti-E microscope body with an Andor Zyla 4.2 PLUS sCMOS camera and an XY linear encoder/piezo Z stage (MP-P500-XYZ-ASI-NKTI) was used. The laser lines used for 405, 488, 561, 594, and 637 nm channels were: 100 mW solid state 405 nm laser; 150 mW solid state 488 nm laser; 150 nM OBIS LS solid state OPSL 561 nm laser; 100 mW OBIS LS 594 nm OPSS laser; and 140 mW OBIS LX solid state 637 nm laser. The respective emission filters were: 450/50; 525/50; 582/15; 631/36; and 676/37. The 405, 488, 561, and 637 nm channels used the dichroic mirror Andor CR-DFLY-DMQD-01 (405/488/561/640). The 594 nm channel used the dichroic mirror Andor CR-DFLY-DMQD-04 (405-445/514/594/730). All images were collected with a Nikon CFI Apo LWD Lambda S 40XC WI NA 1.15 objective, and used 0.4 μm Z-spacing. XY pixel size was 0.1507 μm. All images were collected in confocal mode, unless explicitly noted above.

## Single-embryo RNA-Seq

Embryos were collected at 2-hour intervals for single-embryo RNA seq using SMART-Seq2 low-input RNA seq ^63^. Each embryo was briefly washed with KSOM, then transferred to a lysis buffer in a well of a 96 well plate. Lysate was then frozen on dry ice and stored at -80°C. Sequencing libraries were prepared following the SMART-Seq2 protocol. Libraries were sequenced on an Illumina NextSeq 550 (75 bp, paired-end).

## Feature extraction and data co-registration

The first analysis step was the identification of specific features for each data modality (*in situ* sequencing, immunofluorescence, and mechanics), followed by co-registration of these measurements for the same embryo. This section focuses on embryos characterized with immunofluorescence and *in situ* sequencing, with a subset characterized with mechanical measurements. Code for analyses can be found in: https://github.com/ehabibi1/UTMM.

### Pre-processing

Pre-processing of *in situ* sequencing image data was performed similar to field-of-view image processing in IGS ^22^. Imaris format images were converted to multicolor 3D TIFF stacks (https://github.com/OakesLab/ims-to-tif-converter). Embryo image boundaries (upper left hand corner, length, and width) were manually identified in FIJI for each embryo in each round of imaging. *In situ* sequencing image data (encompassing 7 rounds of *in situ* sequencing, followed by the universal rolony detection hybridization) were then pre-processed as follows. For each embryo, the ROIs corresponding to the embryo boundaries were cropped from multicolor 3D TIFF stacks (which may contain multiple embryos in one FOV). Individual cropped ROIs for each cycle of sequencing were then co-registered to the rolony detection hybridization round (as the reference coordinate system) using normalized cross-correlation on maximum intensity XY and XZ projections of the DAPI channel image stacks to calculate x, y, and z offsets. Calculated offsets were applied to all image channels, and the resulting images, separated into individual 3D TIFF stacks for each channel, were saved.

### Co-registration of morphology and *in situ* sequencing to reference coordinate frame

Morphology (antibody staining) images were co-registered to the rolony detection hybridization round in a similar way. Morphology 3D TIFF stacks were cropped using manually identified borders as before. Three different approaches, described below, were used to co-register the DAPI image from the morphology round to the DAPI image of the rolony detection round, used as the reference coordinate system.

Transformations were applied to all channels and saved as individual single-color TIFF stacks. The best-performing approach, as evaluated by visual inspection, was selected for subsequent processing. The registration approaches are: (1) normalized cross-correlation on maximum intensity XY and YZ projections, as above; (2) phase-correlation on maximum intensity XY projections; and (3) computing a 3D affine transformation.

### Cell segmentation

Morphology images were pre-processed for cell segmentation as follows. A combined p-Ezrin and E-cadherin/Beta-catenin 3D TIFF stack was constructed from the individual p-Ezrin (Ezr) and E-cadherin/Beta-catenin (Ecad) image stacks. First, Ezr and Ecad were median-value subtracted, followed by scaling by dividing by the 99^th^ percentile value. The scaled Ezr and Ecad images were then added, multiplied by 2^16^, and saved as 16-bit TIFF combined images (EcadEzr).

3D Membrane Morphological Segmentation (3DMMS, ^86^) was then used to segment cells based on the EcadEzr combined images, along with DAPI images. Segmentation masks were exported, and examined with 3D Slicer ^87^ for any segmentation artifacts. If needed, masks were manually corrected with 3D Slicer. Segmented cells corresponding to the polar body were deleted from the mask. The final masks were then saved as label matrices.

### *In situ* sequencing – base calling

*In situ* sequencing data was processed following the approach from IGS and code provided herewith using default parameters unless otherwise specified ^22^. Briefly, after initial registration (performed in pre-processing), quantile normalization on each of the image stacks was performed, followed by deconvolution using a high-pass Gaussian filter, and 3D peak finding, localizing each peak to a voxel. For each embryo, thresholds for each color channel (for all rounds of sequencing) were then automatically identified from the distribution of peak intensities using an elbow-finding approach. Peaks exceeding threshold values were retained and expanded to a 5 x 5 x 5 voxel volume (“spots” or “reads”). This volume was integrated for each color channel and sequencing round and quantile normalization was performed on the integrated intensities. Purity values for each spot for each round of sequencing were then calculated as: *purity_n* = max(*c_i*)^2^/(sum_*i*(*c_i*^2^), where *purity_n* is the purity score for the *n*-th round of sequencing for the current spot, and *c_i* where *i* = {1, 2, 3, 4} is the integrated intensity for the *i*-th color in the spot volume in the *n*-th round of sequencing. A purity score for the read was then computed as the number of sequencing cycles for which *purity_n* > 0.7, along with a consensus call for each base (largest value, most likely prediction). Reads with total purity score >= 6 were selected for barcode matching. Barcode assignments were made if the consensus sequence matched to a barcode in the library within a Hamming distance of 1. After processing all spots, spots for the same gene within a Manhattan distance of 4 voxels were combined into one read. The cell segmentation mask for that embryo was subsequently loaded, and used to assign reads to cells based on the center position of each read. Data matrices, containing transcript identity, XYZ coordinates, and cell assignment were then written out, as were cell x gene matrices. Downstream analysis was done using the Seurat package (version 4.3.0, ^88^) to generate the sparse count matrix (see Data normalization section below).

### Extraction of morphological features

Morphology masks were converted into meshes using PyVista (https://pyvista.org)[. Meshes were smoothed and Ezrin signals were mapped to the surface of the mesh and used to extract 3D geometric features using PyRadiomics ^44^. These features consist of:

### Cell shape Features

**Cell Surface Area**: Total outer surface area of the 3D cell.

**Cell Volume**: Total volume occupied by the 3D cell.

**Cell Convex Hull Volume**: Volume of the smallest convex shape that encloses the entire cell.

**Cell Convexity**: A measure of the convexity or concavity of the cell structure, defined as the ratio between the volume of the cell and of its convex hull.

**Cell Center**: Geometric center or centroid of the 3D cell.

**Cell B Sphere Center**: Center of the smallest sphere that can enclose the entire cell (bounding sphere).

**Cell B Sphere Radius**: Radius of the smallest bounding sphere.

**Cell Unit Surface Area**: Surface area of the cell normalized by its volume.

**Cell Exposed Surface Area**: Cell surface area that is exposed to the surrounding environment.

**Proportion Cell Exposed SurfaceArea**: Fraction of the cell’s surface area that is exposed to its environment.

**Cell Major Axis**: Longest axis of the cell.

**Cell Minor Axis**: Axis perpendicular to the major axis but not the least.

**Cell Least Axis**: Shortest axis of the cell.

**Cell Elongation** (Cell stretching): Ratio between the major and minor axes.

**Cell Flatness**: Ratio between the least and major axes.

**Cell Sphericity1 & 2**: Measure of the roundedness of the cell shape relative to a perfect sphere,

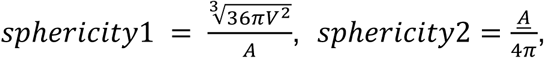

Where *A*, *V*, and *A* are the surface area and volume of the cell, and the surface area of the cell which has been normalized to have unit volume, respectively.

**Cell Compactness1 & 2**

Measure of how compact the cell is relative to a perfect sphere (most compact),

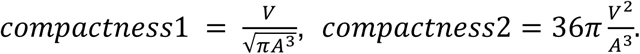

**Cell Spherical Disproportion**

The extent of the cell shape deviation from a perfect sphere. Ratio of the surface area of the cell to the surface area of a sphere with the same volume as the cell,

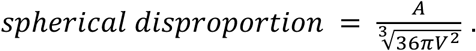

### Curvature Features

#### Curvature Energy

Amount of effort required to bend the cell (indicating its resistance to deformation).

#### Curvature Entropy

Randomness or complexity of the curvature distribution of the cell.

#### Curvature Minimum to Curvature Uniformity

Various statistical measures (e.g., mean, median, max, min, percentile values, variance) of the distribution of curvature across the cell’s surface.

### Textural Features

Ezrin attributes (Ezrin Energy to Ezrin Strength) of the spatial distribution and relationships of pixel intensities within the cell, such as:

**Ezrin Energy**: Sum of squared intensities.

**Ezrin Entropy**: Randomness in intensity value distribution.

**Ezrin Max & Min**: Maximum and minimum intensity values, respectively.

**Ezrin Variance & Std**: Spread of the intensity values.

**Ezrin Skew & Kurtosis**: Shape of the intensity distribution.

**Ezrin Contrast**: Difference between the highest and lowest pixel intensities in a given region.

**Ezrin Correlation**: Given the statistics of how many times the combination of Ezrin intensity levels *i* and j appear in two voxels within the cell, this correlation measures the linear dependency of intensity level values.

**Ezrin Sum Entropy**: Randomness in the sum of pixel values in local regions.

**Ezrin Gray Level NonUniformity**: Variability in gray-level intensity values.

**Ezrin Zone Variance**: Variance in a defined zone of pixel values.

**Ezrin Busyness & Ezrin Coarseness**: Level of detail or granularity in texture. Coarseness measures the average difference, in intensity level, between a center voxel and the average intensity of its 3D neighborhood, in the cell. Busyness is a measure of the average change of intensity between a voxel and a neighbor.

**Ezrin Complexity**: Complexity in the spatial arrangement of pixel intensities. Complexity is high when the space has many rapid changes in Ezrin intensity values.

**Ezrin Strength**: Magnitude of pixel intensities.

### Live-embryo passive microrheology measurements processing

Particle tracking was performed on the raw videos using a customized MATLAB code adapted from ^89^ for batch processing. The two dimensional Cartesian coordinates of tracked mitochondria were recorded and linked into trajectories as a function of time *t*, *r*(*t*). For each mitochondrion, the *RMSD*(*τ*) as a function of lag time *τ* was calculated from

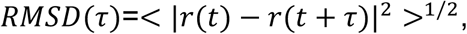

where <> is an average over time *t*. A representative plot of *RMSD*(*τ*) is shown in **fig. S9A**. As expected, *RMSD*(*τ*) shows a short-time plateau, indicating thermal equilibrium at high frequencies. We use RMSD_0_ =*RMSD*(*τ* = 12 *ms*) at the highest imaging frequency to reflect the plateau value of *RMSD*(*τ*), which is proportional to mechanical compliance at thermal equilibrium. The particle tracking yielded a table with RMSD_0_ and positional coordinates for each individual mitochondrion. This pipeline was also used for analyzing microbead motion.

### Optical-tweezers active microrheology

A 1,064-nm laser was used to move the endocytosed 0.5 μm-diameter particles. The optical trap was calibrated in the media. Small amplitude oscillation at a magnitude of 0.2 μm and frequency of 1 Hz was performed at different stages. A complex stiffness k was measured from the force and displacement of the particle. The complex stiffness k was converted to the complex modulus through the Stokes-Einstein relation *G* = *k*/3*πa*, and the magnitude of the complex shear modulus |G| is shown in **Fig. 3C**.

## Electroporation

To selectively label adult neural stem cells (NSCs), we used a combinatory approach and co-electroporated mice with the GFAP-GCaMP6f (or GFAP-GFP) and Prom-mCherry plasmids at P0-P1, as previously described ^56–58,74^. Coincident activity of these GFAP and Prominin (Prom) promoters defines NSCs. For electroporation, P0-P1 CD1 pups were anesthetized by hypothermia and placed in a stereotactic frame. The pups received a unilateral 1-μL injection of both plasmids (4-5 μg/μL) into the lateral ventricle using the following coordinates (with respect to the lambda): anterior-posterior (AP) 1.8 mm, medio-lateral (ML) 0.8 mm, and dorso-ventral (DV) 1.6 mm. We added 1% Fast-Green dye to the plasmid solution to confirm that the plasmids were correctly injected into the lateral ventricle. After the injection, an electric field was rapidly applied to the pups (five 50-ms 100-mV pulses at 950-ms intervals) using an electrode coated with conductive gel and placed on the pup’s bone surface. Electroporated mice were used 2-3 months after. CMV-GCaMP6s (#40753, Addgene) was kindly provided by Dr. Kim & GENIE Project, Janelia Research Campus^90^, and PU6-BbsI-CBh-Cas9-T2A-mCherry (#64324, Addgene) was kindly provided by Dr. Kuehn, Berlin Institute of Health^90,91^. GFP or GCaMP6s was subcloned under the GFAP promoter, while mCherry, derived from the PU6-BbsI-CBh-Cas9-T2A-mCherry plasmid, was subcloned under the Prom promoter.

## Time-lapse imaging in acute brain slices

Acute brain slices were prepared as described previously^92^. Briefly, adult electroporated mice were deeply anesthetized with an intraperitoneal injection of a mixture of ketamine (100 mg/mL) and xylazine (10 mg/mL) and then perfused intracardially with ice-cold modified oxygenated artificial cerebrospinal fluid (ACSF) containing (in mM): 210.3 sucrose, 3 KCl, 2 CaCl2·2H2O, 1.3 MgCl2·6H2O, 26 NaHCO3, 1.25 NaH2PO4·H2O, and 20 glucose. Brains were quickly collected, and 250-μm-thick sagittal slices were cut on a vibratome (VT1000S; Leica). Slices were kept at 37°C in ACSF containing (in mM): 125 NaCl, 3 KCl, 2 CaCl2·2H2O, 1.3 MgCl2·6H2O, 26 NaHCO3, 1.25 NaH2PO4·H2O, and 20 glucose, under continuous oxygenation for no more than 6-8 h.

Before transferring the acute brain sections to the imaging chamber, they were incubated for 1 h with 100 nM of LysoTracker™ DeepRed (Thermo Fisher). Imaging of the fluorescently labeled lysosomes (LysoTracker™) was performed using an upright spinning disk confocal microscope (Nikon) equipped with a 25x water immersion objective (Nikon, NA=1.1) and a Yokogawa CSU-X1 disk. GFP, mCherry, and LysoTracker™ DeepRed were excited by 488-, 561-, and 640-nm solid-state lasers, respectively. The emission was captured using a Prime BSI-Express camera after passing through the appropriate emission and dichroic filters. Labeled lysosomes were imaged in NSCs, identified by the co-expression of GFP and mCherry fluorescence, for 30 s in a single optical section with 2×2 binning, and at an acquisition rate of 75 frames per second.

## Post hoc identification of imaged cells

To identify the imaged cells, 250-μm-thick acute brain sections were fixed with 4% paraformaldehyde (PFA) overnight at 4°C. The acute brain slices were first incubated with ice-cold methanol for 30 min at -30°C, followed by the incubation with ice-cold acetone for 30 min at -30°C. The brain slices were blocked with 4% milk and 0.5% Triton X-100 for 3 h at room temperature and then incubated overnight at 4°C with chicken anti-GFP (1:1000, Aves, *RRID: AB_10000240*), rat anti-mCherry (1:1000, Thermo Fisher, *RRID: AB_2536611*), and rabbit anti-Ki67 (1:1000, Abcam, *RRID: AB_443209*) diluted in 4% milk and 0.5% Triton X-100. The sections were then incubated with the following secondary antibodies diluted in PBS for 3 h at room temperature: Alexa Fluor 488 anti-chicken (1:1000, Thermo Fisher, *RRID: AB_2534096*), Alexa Fluor 568 anti-rat (1:1000, Thermo Fisher, *RRID: AB_2534121*), and Alexa Fluor 633 anti-rabbit (1:1000, Thermo Fisher, *RRID: AB_2535731*). Images were acquired using an upright spinning disk confocal microscope (Nikon) equipped with a 10x air objective (Nikon, NA=0.45) and a Yokogawa CSU-X1 disk.

### Pipeline for co-registration of live-embryo mechanical measurements to fixed-embryo imaging

A semi-automated pipeline was developed to co-register live-embryo and fixed-embryo images in order to map mechanical to transcriptome measurements. These image processing steps were only used to generate the registration transformation, and not for other data analyses.

The main steps in the pipeline consisted of:

For image contrast enhancement, the contrast-fixed and contrast-live functions in ImageJ were respectively used to process fixed and live images to improve clarity and contrast.

These enhanced images served as inputs for the FijiYama plugin in ImageJ, with the output from contrast-fixed as the fixed reference image and the output from contrast-live as the moving image. Manual registration was then conducted using FijiYama’s “Manual Registration - Rigid (no deformations)” function, followed by automatic registration, using the “Automatic registration -> Rigid (no deformations) -> Block-Matching” functionality in FijiYama for further refinement.

Next, relevant spatial coordinates were appended to registered images using add-series-coords function in ImageJ. The register-mechanics function was used to align specific image patches against a reference series. Alignments were user-verified visually and interactively with the align-against-series function. Finally, the write-mecho-locations function documented the identified mechanical locations from the processed and aligned images.

After co-registration, the morphology mask was applied to the mitochondrial RMSD measurements to associate measurements with specific cells.

## Integrative analysis

The second phase of analysis focused on analyzing the features identified previously, along with integrative analysis across features.

## Spatial single-cell analysis of *in situ* sequencing

### Quality control and data normalization

Cells were filtered to retain only higher-quality cells, specifically those with read counts between 100 and 5000 (100 < nCount_RNA < 5000) and feature counts greater than 50 (nFeature_RNA > 50). SCTransform within Seurat was used to normalize the data and perform batch correction, particularly addressing the variables ’plate’ and ‘well’ (vars.to.regress= c(”nCount_RNA”, “nFeature_RNA”, ‘plate’, ‘well’)).

### Dimensionality reduction and clustering

Principal Component Analysis (PCA) was performed on the scaled data using the RunPCA function in Seurat (version 4.3.0). The top 15 principal components (PCs) were retained and a *k*-nearest neighbors graph was constructed based on the Euclidean distance between profiles in the PC space, further refined by shared overlap in local neighborhoods (FindNeighbors, dims = 1:15). Cell profiles were then clustered with the Louvain algorithm (implemented in Seurat) with the modularity function, setting the resolution to 1. Uniform Manifold Approximation and Projection (UMAP) implemented in Seurat with default parameters was used for visualization.

### Differential gene expression analysis

The FindAllMarkers function in Seurat (version 4.3.0, a Wilcoxon Rank Sum test with a Bonferroni correction for multiple testing (adjusted P<0.05)) was used to identify differentially expressed genes with the following parameters: logfc.threshold = 0.25, test.use = “wilcox”, slot = “data”, min.pct = 0.1. Only genes detected in at least 25% of the cells within a cluster and whose average expression was at least 0.25-fold different (log-scale) against the remaining cells were considered for testing.

### Pseudotime analysis

The destiny package (version 3.9.0, ^93^) was used to calculate pseudotime with knn=25, sigma = “local”, n_eigs = 20.

### Non-Negative Matrix Factorization (NMF) for gene programs

NMF (RunNMF, STutility version 0.1.0 implemented in Seurat version 4.3.0) was used to identify gene programs with number of factors = 15. Gene programs were annotated by the top 15 contributing genes. Gene programs were assigned to individual cells with spectral clustering (kernlab version 0.9-32, kernel_function <- “rbfdot”, sigma <-0.1 and k_neighbors <-10).

### Linear Discriminant Analysis (LDA)

LDA was performed with the scikit-learn package (version 1.2.2) in Python. LDA, with two components, was applied on the 64 morphological features with the cell states: ICM-1, ICM-2, pmTE, and TE-1as response variables.

### Cell Alignment and Polarization Analysis

To investigate the spatial polarization of TE-2 and TE-1 cells across embryos, we implemented an algorithm that aligns and aggregates cell distributions based on a unit vector that maximally separates the two populations. This approach ensures consistent alignment of spatial features across embryos.

**Step 1**: Relative Position Calculation

For each cell within an embryo, compute the relative position of the cell to the embryo’s center:

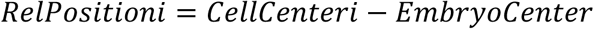

**Step 2**: Embryo Normalization

To account for size differences across embryos, normalize the relative positions within each embryo by the maximum Euclidean norm of any cell in the embryo:

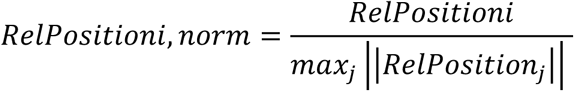

**Step 3**: Population Separation Optimization

To find the unit vector *v* that maximally separates the TE-2 and TE-1populations, solve the following optimization problem:

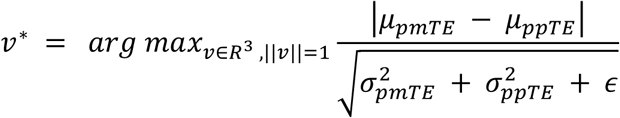

where: *μ_pmTE_* and *μ_ppTE_* are the means of the projections of TE-2 and TE-1 cells along *v*, respectively. 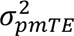 and 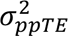 are the variances of the projections. ∈ is a small regularization term to prevent division by zero.

**Step 4**: Reorientation for Consistent Alignment

To ensure consistent alignment across embryos, the orientation of *v* is flipped if necessary, such that the TE-1 population consistently aligns with positive projections along *v*.

**Step 5**: Aggregation and Visualization

After alignment, the projections of all cells along *v* are aggregated across embryos to analyze spatial polarization and distribution. The resulting data is visualized as density plots and scatter plots.

### Volumetric compression RNA-seq analysis

Raw sequencing reads was converted to fastq files using bcl2fastq (version v2.20.0). Reads were aligned to the reference mouse genome GRCm38 and Ensembl gene annotation (release 102). Reads were aligned with Bowtie (version 0.12.9) and bam2hits and mmseq package ^94^ were used to generate gene-by-embryo matrices.

Gene-by-embryo matrices were grouped into control (KSOM media) and treated (KSOM + 2% PEG 300) sets, and further partitioned into early (4-6 hours) and late (12-15 hours) response embryos. Differential expression analysis was performed with DESeq2 version 1.26.0 ^95^, followed by enrichment of gene annotations, focusing on pathways within the Kyoto Encyclopedia of Genes and Genomes (KEGG) database (Genetic Information Processing, Environmental Information Processing, Cellular Processes, and other relevant biological functions), with Fisher’s exact test, and the Benjamini-Hochberg False Discovery Rate (FDR) method.

### Time-lapse imaging and volumetric analysis of compressed embryos

Early 4-cell-stage embryos were stained with 100 nM CellTracker for 30 min before imaging and subsequently maintained in a culture medium containing 10 nM CellTracker throughout the experiment. Control and volumetrically compressed embryos (N = 7 embryos per condition) were imaged for 5 h on a Leica SP8 point-scanning confocal microscope under controlled culture conditions at 37 °C and 5% CO₂. Images were acquired every 30 min using 1% laser power. For volumetric analysis, the whole embryo was segmented at each time point based on the CellTracker fluorescence signal. A three-dimensional mask encompassing the entire embryo was generated from the CellTracker-positive volume, and total embryo volume was calculated from this mask. For each embryo, the volume measured at each time point was normalized to its volume at the beginning of the imaging period.

## List of Supplemental Materials

**Supplemental Data 1:** Branching trees showing the expression of all 150 genes from scRNAseq data used for 3D targeted *in situ* sequencing.

**Supplemental Table 1:** Details of the probes used for 3D targeted *in situ* sequencing: gene symbols, accession numbers, and barcodes information.

**Supplemental Table 2:** Details of the probes used for 3D targeted *in situ* sequencing: probe sequences.

**Supplemental Table 3**: Details of the probes used for 3D targeted *in situ* sequencing: oligo sequences.

**Supplemental Table 4-5:** Experimental metadata and summary statistics of collected embryos

**Supplemental Table 6:** 3D targeted *in situ* sequencing and multiplex HCR comparison data

**Supplemental Table 7:** Differentially expressed genes between early control (4cell) vs. PEG (4cell).

**Supplemental Table 8:** Differentially expressed genes between early control (4cell) vs. late control (8cell).

**Movie S1:** Representative spontaneous cytoplasmic motion in a cell at morula stage visualized in a raw video using MitoTracker (orange), LysoTracker (red) and endocytosed fluorescent tracer particles (green). Trajectories for each particle are overlaid in lighter shades of the same colors used for their markers (orange for MitoTracker, red for LysoTracker, and green for beads), showing their motion over time

**Movie S2:** Representative particle tracking results overlaid with raw videos of one embryo. Each grid is a raw video of MitoTracker. The red circles mark the tracked mitochondria that are used to calculate RMSD.

**Movie S3:** Representative spontaneous cytoplasmic motion in a 4 cell embryo visualized with endocytosed fluorescent tracer particles. In this work we only focus on the short time-scale movement in thermodynamic equilibrium, while directed active movement can be observed at long time scales.

**Movie S4:** Representative spontaneous cytoplasmic motion in a 8 cell embryo (compacted) visualized with endocytosed fluorescent tracer particles. In this work we only focus on the short time-scale movement in thermodynamic equilibrium, while directed active movement can be observed at long time scales.

**Movie S5:** Representative spontaneous cytoplasmic motion in a blastocyst stage embryo visualized with endocytosed fluorescent tracer particles. In this work we only focus on the short time-scale movement in thermodynamic equilibrium, while directed active movement can be observed at long time scales.

